# Spatial segregation and aging of metabolic processes underlie phenotypic heterogeneity in mycobacteria

**DOI:** 10.1101/2023.12.01.569614

**Authors:** Celena M. Gwin, Kuldeepkumar R. Gupta, Yao Lu, Lin Shao, E. Hesper Rego

## Abstract

Individual cells within clonal populations of mycobacteria vary in size, growth rate, and antibiotic susceptibility. Heterogeneity is, in part, determined by LamA, a protein found exclusively in mycobacteria. LamA localizes to sites of new cell wall synthesis where it recruits proteins important for polar growth and establishing asymmetry. Here, we report that in addition to this function, LamA interacts with complexes involved in oxidative phosphorylation (OXPHOS) at a subcellular location distinct from cell wall synthesis. Importantly, heterogeneity depends on a unique extension of the mycobacterial ATP synthase, and LamA mediates the coupling between ATP production and cell growth in single cells. Strikingly, as single cells age, concentrations of proteins important for oxidative phosphorylation become less abundant, and older cells rely less on oxidative phosphorylation for growth. Together, our data reveal that central metabolism is spatially organized within a single mycobacterium and varies within a genetically identical population of mycobacteria. Designing therapeutic regimens to account for this heterogeneity may help to treat mycobacterial infections faster and more completely.

## Introduction

For a bacterial infection to linger after antibiotic treatment, only a few bacteria need to remain. What is different about surviving bacteria, and how do these differences arise? Despite extensive research, no single model has emerged explaining this phenomenon, and it is likely that the answers to these questions will vary depending on the bacterial species. In the case of model bacteria, like *Escherichia coli*, much of the focus has been on stochastic mechanisms that underlie cells switching into rare drug-tolerant states (*1, 2*). However, stochasticity is just one way of generating diversity, and other mechanisms of heterogeneity are more deterministic in nature. For example, clonal populations of mycobacteria, a genus that includes the human pathogen *Mycobacterium tuberculosis*, exhibit more variability than model bacterial species (*3*). At least some of the heterogeneity is created every time a mycobacterium divides when it produces two cells with different sizes, growth rates, and susceptibilities to antibiotics (*4*). Importantly, heterogeneity is hard-coded in the genome, as deletion of a single gene unique to mycobacteria – *lamA* – collapses morphological heterogeneity and leads to fast and uniform killing by several antibiotics (*5*). LamA localizes to sites of new cell wall synthesis and recruits proteins important for polar growth (*6*). However, many bacterial species divide asymmetrically but do not exhibit as much heterogeneity as mycobacteria, suggesting that asymmetric division is only one factor responsible for creating a heterogeneous population (*7*). Here, we sought to understand the mechanism by which LamA creates heterogeneity in growth.

## Results

### A conserved tyrosine regulates the localization and function of LamA

The predicted structure of LamA includes two regions of high disorder separated by a single transmembrane domain (**Fig. 1A**). The carboxy terminus of the protein is predicted to encode an MmpS domain. Proteins with these domains are often encoded by operons that include *mmpL* genes, but LamA’s operon partner is a gene of unknown function. The amino terminus of LamA is highly acidic, with nearly 20% of the residues in this region being either aspartic or glutamic acid. Additionally, several whole-cell proteomic studies have mapped phosphorylation events at serine, threonine, and tyrosine residues in this region (*8-13*). We performed a multiple sequence alignment of LamA proteins found throughout the mycobacterial genus and observed that, despite the predicted disorder of this region, several tyrosine and serine residues were highly conserved (**Fig. 1B**). To test the function of these residues, we created phage-integrating plasmids carrying alanine mutations at each of these sites, transformed these into *ΔlamA M. smegmatis*, and imaged the resulting strains by phase contrast microscopy. To quantify the morphology of single cells, we trained the U-Net machine-learning software package to detect and segment single mycobacterial cells by phase contrast (*14, 15*). Using this approach, we found that one mutant, a tyrosine mutated to an alanine at position 50, resulted in cells that were slightly larger than the other mutants and wild type (**Fig. 1C**). Increased cell size was due to an increase in cell length rather than width (**Fig. S1**). Importantly, tyrosine 50 is found to be phosphorylated in several whole-cell proteomic studies (*9-11*). To confirm that this mutation is important at the native locus, we used marker-less single-strained recombineering (*16*) to recode this tyrosine to an alanine at the native locus and performed timelapse microscopy. Compared to wild type cells, a subpopulation of cells expressing LamA_Y50A_ was indeed larger, driving the increased size of cells across the population (**Fig. S2**). This also led to more heterogeneity in cell size at the time of division; however, it did not change the asymmetry at division (**Fig. 1D, Fig S2**). Both phenotypes suggested that LamA_Y50A_ functions similarly to LamA_WT_ or may be a gain-of-function mutant.

**Figure 1.**
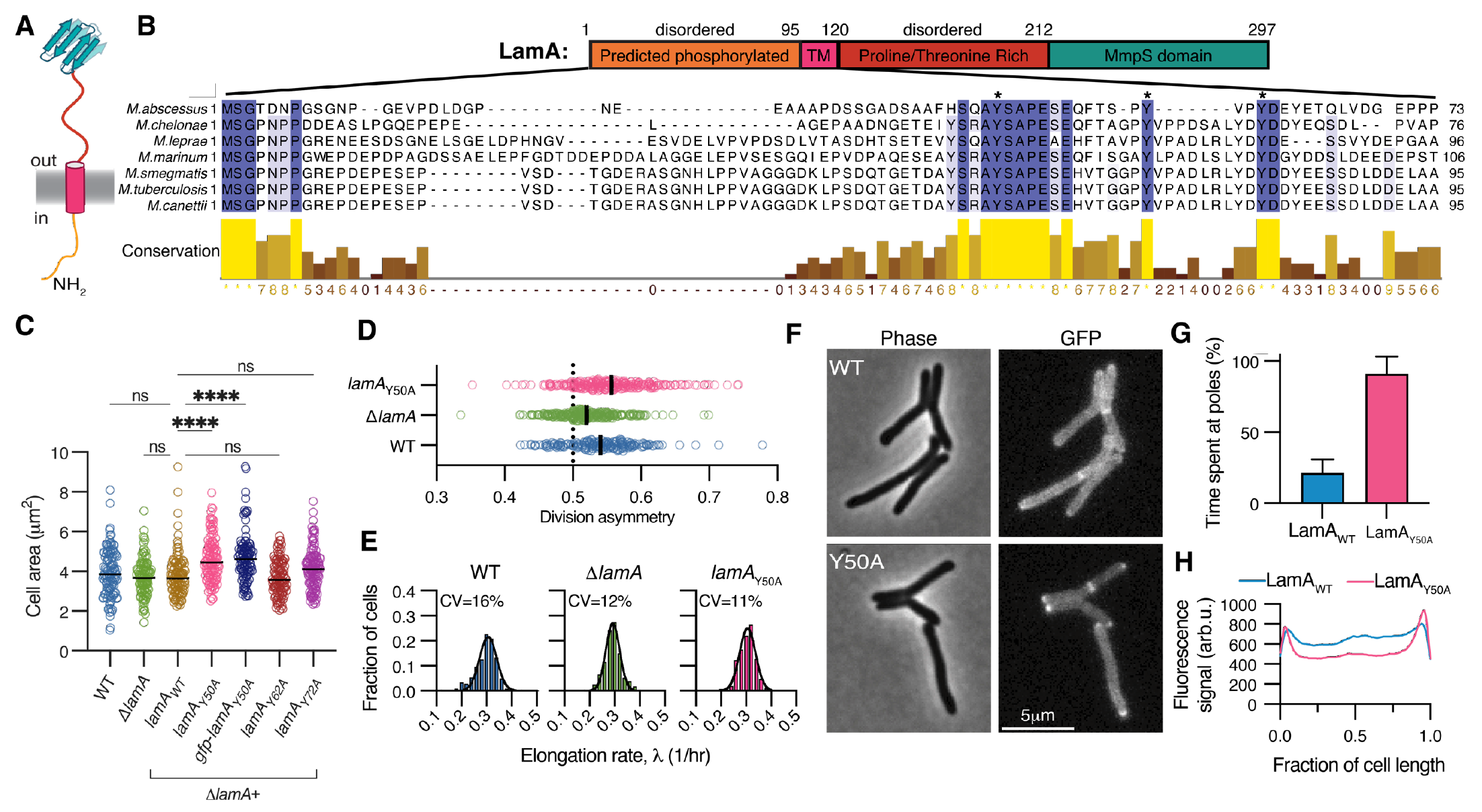
A conserved phosphorylated tyrosine is important for LamA function and localization. Schematic of the predicted structure of LamA. **(B)** A multiple sequence alignment of the N-terminal uence of LamA. Stars indicate residues mutated in panel C. **(C)** Cell size of WT, Δ*lamA*, or cells ressing the indicated *lamA* allele in Δ*lamA*. (dark black lines indicate medians; n=100-125 cells for strains; ****p<0.0001 by one-way ANOVA comparing means to *lamA*_WT_ corrected for multiple mparisons). **(D)** Asymmetry in the daughter cell size at the time of division, measured by the old pole ghter as a fraction of the total size of both daughters. The dotted line marks symmetric division. 195, 236, 223 daughter cell pairs for WT, Δ*lamA*, and *lamA*_*Y50A*_, respectively). **(E)** The elongation rate, f the indicated strains. (CV = coefficient of variation, n=239, 226, 267 single cells for WT, Δ*lamA*, and *A*_*Y50A*_, respectively). **(F)** Phase contrast and fluorescence images of either msfGFP-LamA or fGFP-LamA_Y50A_ expressed in single copy from the native promoter. **(G)** Fractional dwell time of strains anel F calculated by the amount of time a focus spent at the pole before dissipating, normalized to total time of a fast-interval timelapse (n=15 cells for each strain). **(H)** Intensity profiles were measured oss several cells, normalized to cell length, and averaged (n =140-165 cells for each strain). In panels E, LamA_Y50A_ refers to the strain mutated on the chromosome in an otherwise wild type background. ll others, *lamA*_*Y50A*_ is expressed in single copy in Δ*lamA*.

However, this mutant was deficient in other functions performed by LamA. Specifically, LamA also affects the heterogeneity in single-cell growth rate. Biochemical fluctuations within metabolic pathways result in individual cells growing at slightly different rates centered around a population average (*17*). To compare growth rate distributions between strains, we relied on the observation that single *M. smegmatis* cells grow exponentially and computed an average growth rate, λ = ln(S_d_/S_b_)/ΔT, where S_d_ is the size of the cell at division, S_b_ is the size of the cell at birth, and *Δ*T is the time between birth and division (*18*). Wild type *M. smegmatis* displayed 16% variation in λ, in agreement with prior studies that computed λ by measuring length instead of area (*18, 19*). In contrast, both *ΔlamA* and LamA_Y50A_ cells were less variable than wild type (CV_λ_ = 12% and 11%, respectively) (**Fig. 1E**). Computing growth by other metrics resulted in the same conclusion (**Fig. S3A**). Importantly, expression of LamA_WT_ from an integrative plasmid restored heterogeneity in *ΔlamA* (**Fig. S3B**). Thus, LamA has at least two functions. One establishes asymmetry in cell size at the time of division; the other is dependent on tyrosine 50 and mediates heterogeneity in the growth rate of single cells.

To understand what was different about LamA_Y50A_, we fused msfGFP to either the wild type or mutant variant of LamA at their N-termini and expressed these from the native promoter at a phage integration site in *ΔlamA*. We had previously determined that LamA localized to the septum, but were unable to visualize membrane localization, possibly due to over-expression of the fusion construct (*5*). By expressing msfGFP fusions at native levels, we observed that, in addition to localizing to the septum, msfGFP-LamA_WT_ also localizes along the sides and is occasionally found at the poles (**Fig. 1E**). Timelapse imaging at 5-minute intervals confirmed that msfGFP-LamA_WT_ is highly dynamic between the pole and sides of the cell, at a timescale that is too fast to represent new synthesis and maturation of the fluorescence protein fusion construct (**Fig. 1F-H)**. We next localized msfGFP-LamA_Y50A_ and found its localization to be much less dynamic, with the protein primarily localized to the poles (**Fig. 1F-H**). Interestingly, the human pathogen *M. tuberculosis* divides less asymmetrically than *M. smegmatis* but, like *M. smegmatis*, exhibits a wide variation in single-cell growth rates (*20*). As we have connected LamA to both asymmetric division and growth rate heterogeneity, we wondered if the *M. tuberculosis* LamA variant localized differently than *M. smegmatis* LamA. Indeed, we find the *M. tuberculosis* variant of LamA expressed in *M. smegmatis* is mainly found along the side walls, with less polar localization than we observe with the *M. smegmatis* variant (**Fig. S4**).

Taken together, these data reveal that the functions of LamA are performed at different subcellular sites and that LamA dynamically localizes between these sites, possibly in a phosphorylation-dependent manner. Specifically, we find that a mutant of LamA that cannot be phosphorylated at a conserved residue is locked at the poles and grows more uniformly. This suggests that the mechanisms used to create heterogeneity in growth are performed along the sides of the bacterium. Furthermore, these data suggest that LamA functions at the pole to establish asymmetric polar growth. In fact, we have recently shown that LamA is important for recruiting certain proteins to the poles, which are required to establish asymmetry (*6*).

### LamA precipitates with proteins involved in oxidative phosphorylation, which are excluded from the poles and septa

As LamA is performing different functions at distinct subcellular sites, we hypothesized that it would interact with different proteins depending on its localization. To examine this, we performed a series of immunoprecipitations to find potential protein-protein interaction partners. We created a strain in which the sole copy of LamA was fused to the 3X-FLAG epitope. We fixed these cells with a chemical crosslinker to capture potentially transient interactions, incubated lysates with α-FLAG beads, and identified co-precipitating peptides by mass spectrometry. Precipitated peptides largely fell into two categories of proteins – those associated with cell elongation (MmpL3, PgfA, MurA, and PknA), and those associated with cellular respiration and oxidative phosphorylation (components of the electron transport chain and ATP synthase complex) (**Table S1**). To verify these results, we conducted a reciprocal co-immunoprecipitation with AtpG, the gamma subunit of ATP synthase, since it was highly enriched in multiple biological replicates. For this, we created strains expressing both AtpG-3XFLAG and LamA-strep and found that we could precipitate LamA-strep with α-FLAG beads only when AtpG-3XFLAG was present (**Fig. 2A**).

**Figure 2.**
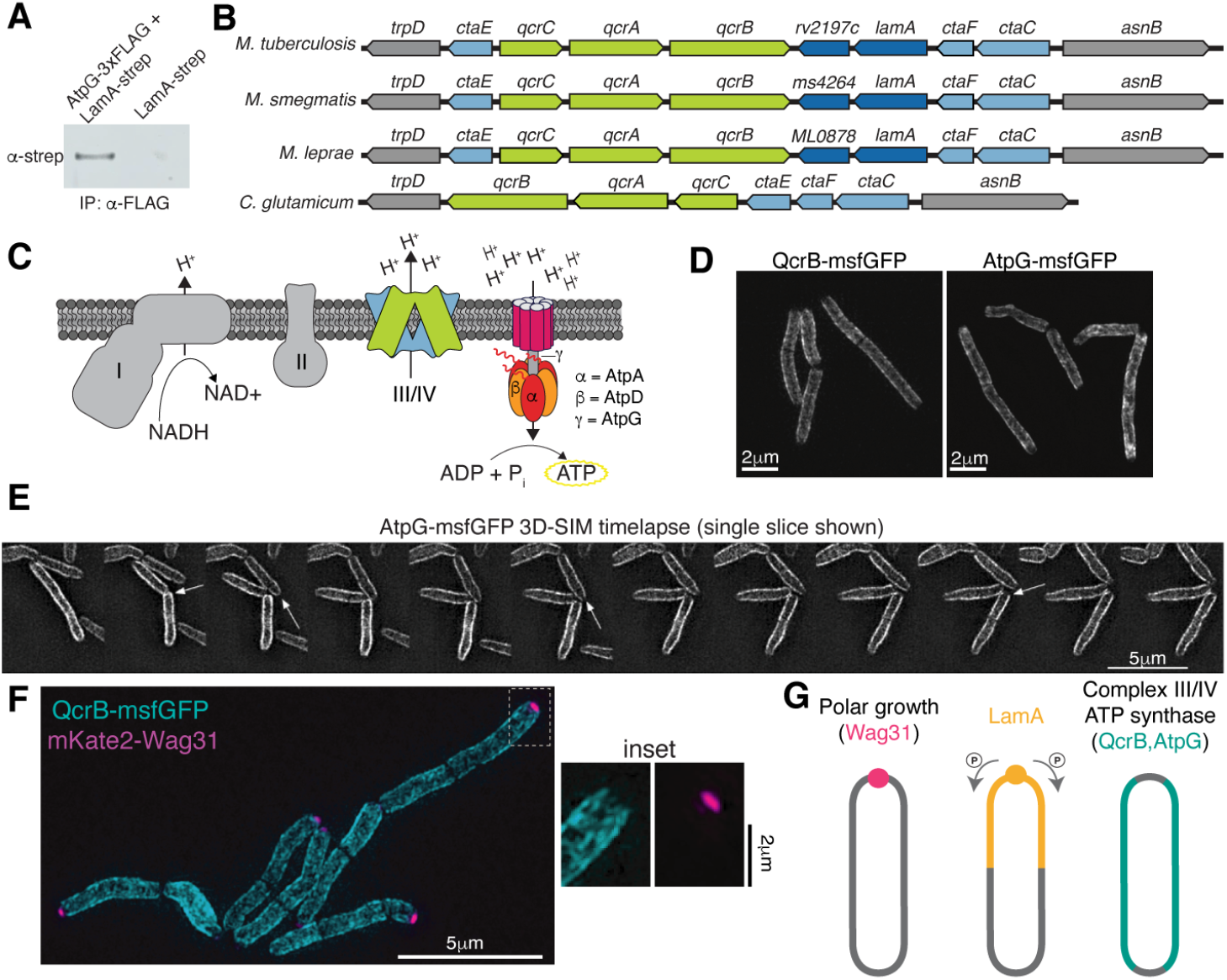
LamA interacts with proteins involved in OXPHOS, which are spatially separated from those used to grow the cell. **(A)** Strains expressing the indicated epitope-tagged proteins were incubated with α-FLAG beads, and the precipitates were probed with an α-strep antibody via Western blot. (**B)** The genomic region surrounding *lamA* in the indicated organisms. **(C)** The proteins in the ETC that correspond to the genes in panel B. **(D)** Live cells expressing the indicated fusions were imaged by 3D-SIM. A maximum projection is shown. **(E)** Cells expressing AtpG-msfGFP were followed over time by 3D-SIM at 15-minute increments. At each time point single slices from the 3D stacks are shown. Arrows indicate AtpG dynamically infiltrating the tips of cells. **(F)** Cells expressing fusions to QcrB and Wag31 were imaged by live-cell 3D-SIM. A maximum projection is shown. **(G)** A model summarizing the localization of LamA and the complexes involved in polar growth and OXPHOS.

Perhaps shedding light on these interactions, the operon containing *lamA* is arranged in the middle of two operons containing many of the genes that encode proteins in the cytochrome III/IV supercomplex, which performs the last step in the electron transport chain (ETC). This genomic synteny is conserved across mycobacterial species (**Fig. 2B,C**). To further investigate the connection between LamA and the ETC, we tracked the localization of QcrB, a subunit of the III/IV supercomplex, by making an in-frame fusion to msfGFP at the chromosomal locus. We also created fusions to AtpG and AtpA, the gamma and alpha subunits of the ATP synthase complex, respectively. As expected, by conventional fluorescence microscopy, the proteins were localized to the plasma membrane (**Fig. S5A**). To visualize the localization of these proteins more closely, we loaded cells into a microfluidic device and performed live-cell three-dimensional structured-illumination microscopy (3D-SIM), a super-resolution technique. As SIM requires collecting several images to reconstruct one super-resolved image, we focused on AtpG and QcrB fusions because these were the brightest. We collected images at single time points and observed that both proteins were excluded from the tips of the cells as well as the septa of dividing cells (**Fig. 2D**). By timelapse 3D-SIM at 15-minute increments, we occasionally saw AtpG-msfGFP infiltrate the tip of the cell but primarily remain localized to the sides of the cell (**Fig. 2E**). To visualize the localization in the context of proteins that direct polar growth, we created a strain that encoded key polar growth scaffolding protein, Wag31, fused to mKate2 in the background of cells expressing QcrB-msfGFP and performed two-color live-cell 3D-SIM (**Fig. 2F,G**). We observed little to no colocalization between these two proteins, showing that the complexes that perform the last steps of oxidative phosphorylation are spatially distinct from those involved in growing the bacterium (**Fig. 2F, Fig. S5B)**. Moreover, as LamA is localized dynamically between the poles/septa and sides, these data also suggest that the interaction between LamA and the electron transport chain is transient and largely confined to the sides, rather than poles, of the bacterium (**Fig. 2G**).

### LamA inhibits oxidative phosphorylation, and reliance on oxidative phosphorylation is associated with uniformity

If LamA is working with the respiratory chain, we reasoned that changes to the membrane potential and cellular ATP levels might reveal if it has a negative or positive regulatory role. We measured membrane potential by quantifying the accumulation of a charged fluorescence molecule, TMRM, by flow cytometry (*21, 22*). As a control, we treated cells with a known protonophore, CCCP, at greater-than-MIC concentrations to depolarize the membrane and observed the expected decrease in signal (**Fig. 3A**). Compared to wild type, *ΔlamA* was slightly depolarized (**Fig. 3A**), supporting the idea that LamA is functioning with members of the electron transport chain. Next, we measured ATP levels using a standard luminescence-based assay and found that ATP levels were slightly higher in *ΔlamA* cells (**Fig. S6A**). Together, these data suggest that LamA may be inhibiting the production of ATP via oxidative phosphorylation.

**Figure 3.**
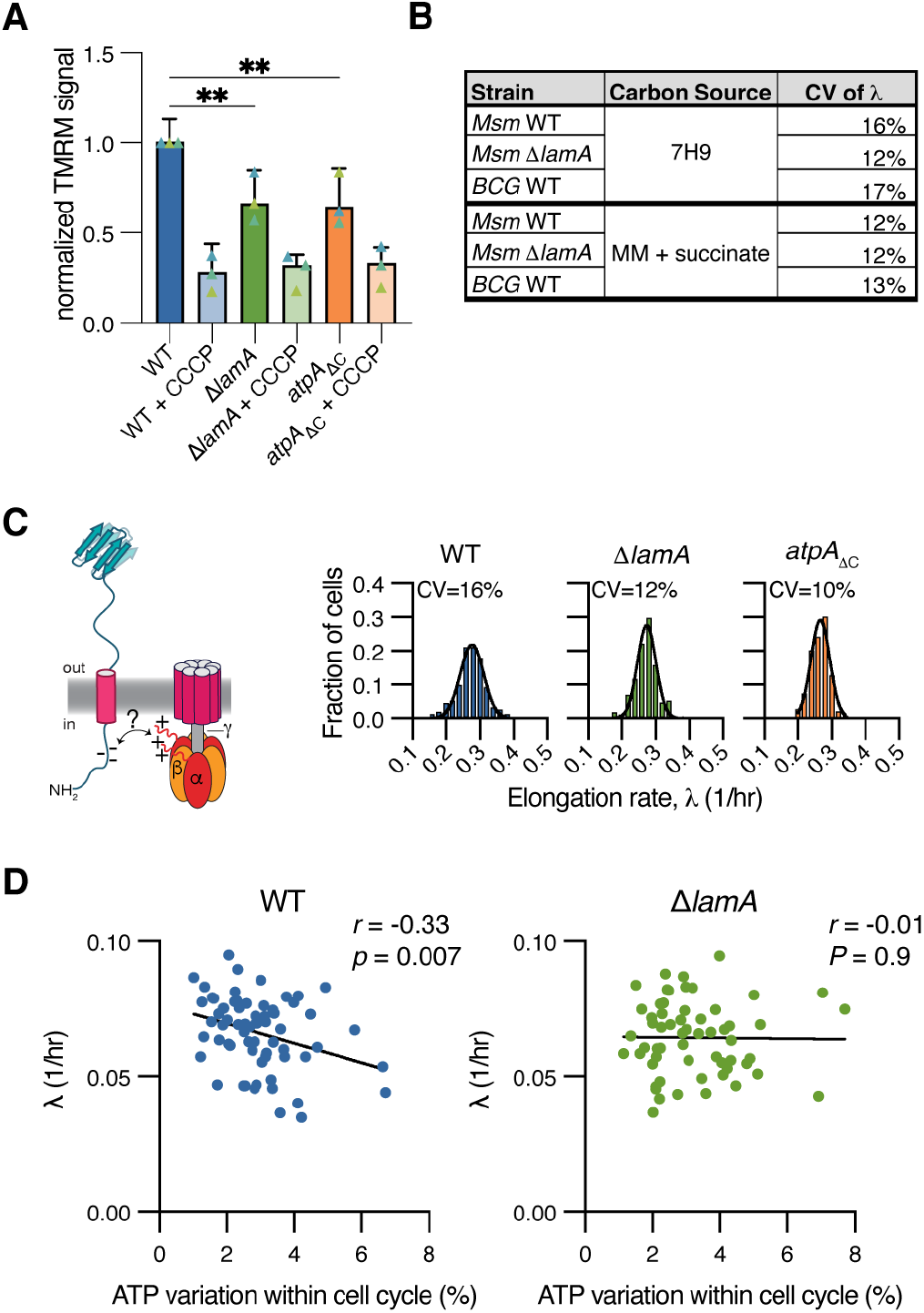
LamA-mediated heterogeneity is tightly linked to ATP production. **(A)** Accumulation of TMRM is measured by flow cytometry. As a control, cells were treated for 15 minutes with 500μM of CCCP. The bars represent the medians of three experiments (each with 3 biological replicates (triangles)) assayed on different days. Error bars are 95% confidence intervals. **p<0.002 by one-way ANOVA. **(B)** The indicated strains were grown in either 7H9 or minimal media (HdB for *Msm* or MMAT for *BCG*) supplied with succinate as the sole carbon source and imaged by phase contrast timelapse microscopy to compute λ for single cells. **(C)** Mycobacteria encode an unusual domain in the alpha subunit of the ATP synthase that prevents hydrolysis. We hypothesize that this positively charged extension interacts with the negatively charged cytoplasmic tail of LamA. Cells missing this extension were imaged over time to measure λ. (n = 152, 172, and 149 cells for WT, *ΔlamA*, and *atpAΔC* cells, respectively). **(D)** WT or *ΔlamA* cells expressing the QUEEN-2m ATP biosensor were imaged over time, and the ratio of the two QUEEN excitation wavelengths was used to obtain relative ATP levels. For each cell, the variation within a cell cycle (as measured by the standard deviation of the QUEEN-2m signal divided by its mean) was compared to the exponential growth rate (n = 67 and 59 complete cell cycles for WT and *ΔlamA*, respectively; lines represent linear regressions; *r* is Pearson’s correlation coefficient; and, *p* is the *p*-value).

Oxidative phosphorylation is essential for mycobacteria, but mycobacteria can also produce ATP in other ways, like substrate-level phosphorylation when grown on sugars. We wondered how generalizable the connection between metabolism and heterogeneity was. Specifically, we asked if cells grown in conditions promoting the exclusive use of oxidative phosphorylation would phenocopy Δ*lamA*. To test this, we grew cells in defined minimal media and supplied succinate as the sole carbon source. Consistent with our hypothesis, wild type *M. smegmatis* grew uniformly in minimal succinate medium, and deletion of *lamA* had no additional effect on heterogeneity (**Fig. 3B**). *M. smegmatis* cells cultured in acetate also grow uniformly (*19*), suggesting this phenomenon is not specific to succinate-grown cells. To understand if this was also true in slow-growing mycobacteria, we followed single *Mycobacterium bovis* BCG cells for approximately four doublings (∼4 days) in both our normal medium (7H9) and minimal medium supplied with succinate. Consistent with our findings in *M. smegmatis*, BCG cells cultured in 7H9 grew more heterogeneously than those fed only succinate (CV_λ_ = 17% and 13%, respectively) (**Fig. 3B**). Together, these data show that cells undergoing more oxidative phosphorylation, either through deletion of *lamA*, or through carbon source availability, grow more uniformly.

### An unusual extension on the ATP synthase alpha subunit mediates heterogeneity

We hypothesized that the interaction between LamA and OXPHOS proteins is important for creating heterogeneity in the growth rate of single cells. The complexes that perform oxidative phosphorylation in mycobacteria are comprised of highly conserved proteins, but there are several unique characteristics specific to mycobacteria and related species (*23*). For instance, in many organisms, ATP synthase can both synthesize and hydrolyze ATP depending on cellular conditions. However, in some actinobacteria, including mycobacteria, the alpha subunit of ATP synthase has a disordered extension (residues V519 – A548) that interacts with the gamma subunit to prevent the enzyme from hydrolyzing ATP (*24*). The ATP synthase operon is duplicated in *M. smegmatis (25)*; therefore, we created a strain in which we deleted the extension in both copies of AtpA (*atpA*_*ΔC*_). Consistent with the known function of this extension to block ATP hydrolysis, *atpA*_*ΔC*_ cells have slightly lower ATP levels (**Fig. S6B)**.

Because the extension on AtpA is highly positively charged, and the N-terminal extension of LamA is negatively charged both by amino acid residues and by multiple phosphorylated residues, we hypothesized that these two proteins might be interacting electrostatically (**Fig. 3C**). To that end, we reasoned that if LamA interacts with this extension to inhibit ATP synthase, a condition we have shown is associated with more heterogeneity, then we should expect that AtpA_ΔC_ cells would phenocopy Δ*lamA* cells with regards to both membrane potential and single-cell growth heterogeneity. To test membrane potential, we used the same TMRM assay as before and found that the depolarization in *atpA*_*ΔC*_ cells was similar in magnitude and direction as that of *ΔlamA* cells (**Fig. 3A**). Next, we analyzed single-cell growth by phase contrast timelapse microscopy, and consistent with our hypothesis, *atpA*_*ΔC*_ cells exhibit less variability in growth than wild type cells (**Fig. 3C**), but not division asymmetry **(Fig. S7)**. Further experiments will be needed to determine if LamA and ATP synthase interact directly, a challenging directive as our data suggests any interaction is likely transient. Nevertheless, we conclude that both LamA and an actinobacteria-specific feature of the ATP synthase are needed to create phenotypic heterogeneity within a genetically identical mycobacterial population.

### LamA mediates the coupling between ATP fluctuations and growth in single cells

Cellular ATP levels are often assumed to be uniformly distributed across a population and stable over the course of a cell cycle. However, fluorescent biosensors that dynamically report on ATP concentrations in single bacterial cells have revealed that ATP levels vary from cell to cell and fluctuate dynamically within a cell cycle (*26, 27*). For single cells within a clonal population, large ATP fluctuations during a cell cycle are associated with slower growth (*28*). While the biological basis for this phenomenon remains unknown, it offers a potential explanation for phenotypic heterogeneity in growth rate. Since we show that LamA is connected to ATP generation through oxidative phosphorylation, we hypothesized that ATP fluctuations would be altered in *ΔlamA* cells. To assay this, we expressed a codon-optimized version of the QUEEN-2m biosensor in *M. smegmatis* cells with and without *lamA (26)*. ATP concentration within bacterial cells is linearly proportional to the ratio of fluorescence excited at 405nm versus 488nm detected by the same emission bandpass (*26*). Thus, we computed pixel-by-pixel ratios of the fluorescence values collected at these wavelengths. As in *E. coli (26)*, single-cell measurements revealed a negative correlation between the amplitude of fluctuation of QUEEN-2m signal (*i.e*. [ATP]) and growth rate in wild type *M. smegmatis* cells. On average, Δ*lamA* cells displayed a similar magnitude of ATP fluctuation, but these fluctuations were not correlated with growth rate (**Fig. 3D**). Together, these data show that LamA mediates the coupling between ATP production and growth rate across a clonal population of *M. smegmatis* cells.

### Mycobacterial single-cell aging is associated with metabolic heterogeneity

After division, rod-shaped cells inherit a new pole formed from the most recent division event and an old pole that was formed during a prior division event. For mycobacteria, this means that cells inherit growing poles of various ages (**Fig. 4A**). Cells with the oldest poles are born larger but grow more slowly and with more variability (*19, 29-31*). As we have connected growth variability to the production of ATP, we wondered how the abundance of OXPHOS components differed as single cells aged. By timelapse microscopy, we observed that the fluorescence intensity of QcrB-msfGFP (a proxy for the cellular concentration of QcrB) was inherited unevenly at division. On average, “old pole” cells inherited a lower concentration of QcrB than their “new pole” siblings. This difference was reduced in Δ*lamA*, with a more uniform inheritance of subunits between sisters (**Fig. 4A**).

**Figure 4.**
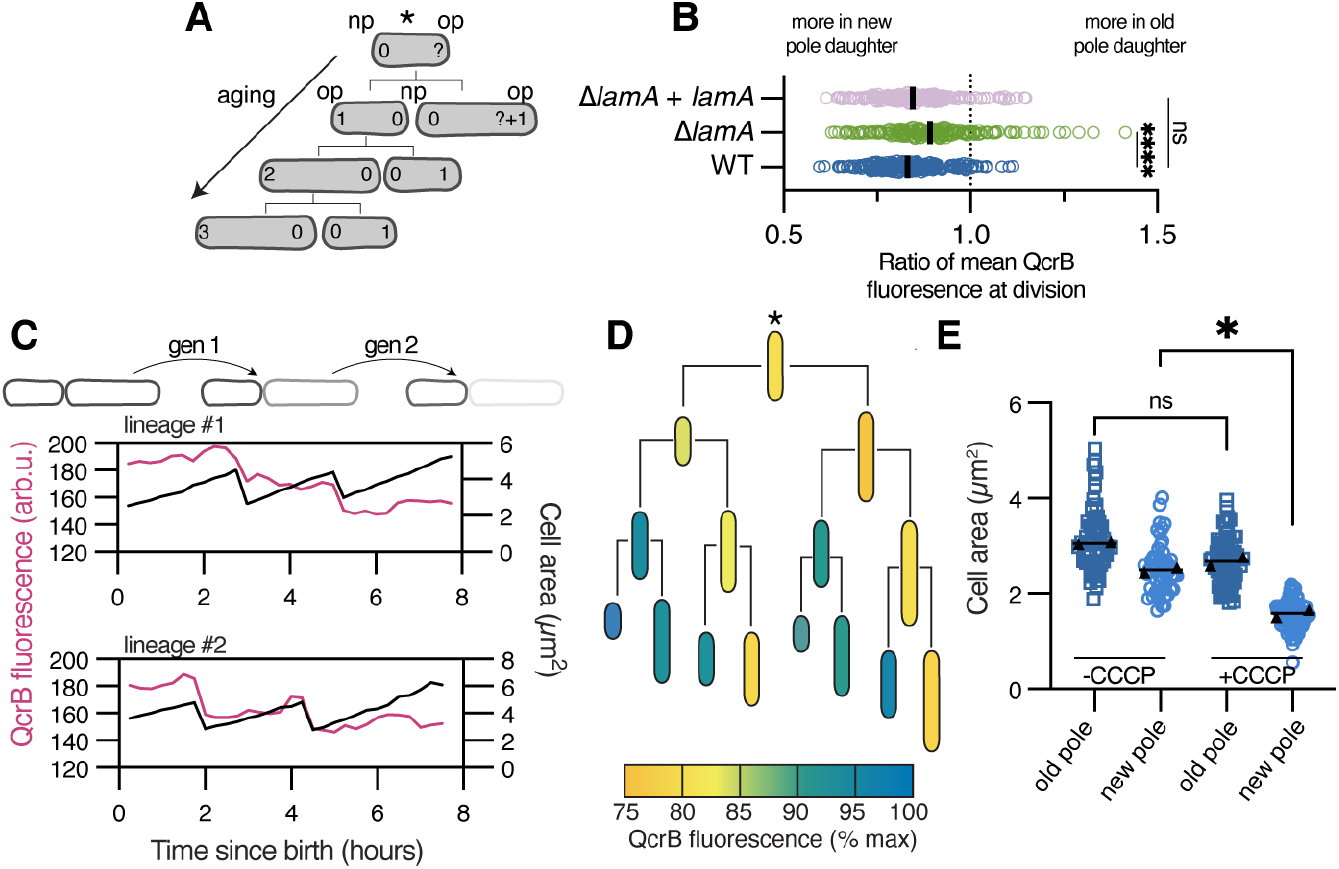
Metabolic aging in single mycobacterial cells. **(A)** Schematic of aging cells and their associated pole ages. **(B)** QcrB-msfGFP fluorescence in the new pole daughter is compared to the fluorescence in the old pole daughter at the time of division in the indicated strains. The dotted line represents equal inheritance. **** p<0.0005 by one-way ANOVA. (n = 187, 191, and 159 sister pairs for WT, λ*lamA*, λ*lamA* +*lamA*, respectively). **(C)** QcrB-msfGFP fluorescence was followed over time in old pole daughters as they age. Two representative lineages are shown. **(D)** QcrB-msfGFP fluorescence over multiple generations, coloring corresponds to percent signal normalized to the maximum (n = 10 lineages). The initial mother cell is of unknown age as indicated by the * in panel A. **(E)** Cells grown in a microfluidic device and treated with 50μM CCCP. Size was measured pre- and post-treatment. (From left to right: n = 43, 40, 64, 47 over two different experiments, indicated by the triangles). P<0.05 by one-way ANOVA comparing the means of the triangles.

We next asked how these differences propagated through the generations in a wild type population. Thus, we identified the youngest cells in our timelapse data and followed them for at least two generations. For many old pole daughters, we observed that fluorescence decreased at time of division and declined steadily over multiple divisions (**Fig. 4C, 4D**). Analyzing multiple lineages showed that QcrB concentration decreased in the old pole daughter cells by approximately 25% after the third division (**Fig. S8A**,**B**). Reconstructing complete lineages revealed that this phenomenon occurred to a greater or lesser degree depending on the identity of the mother cell and was most pronounced in cells with the oldest mothers (**Fig. 4D**). We repeated these measurements with cells expressing AtpG-msfGFP and observed similar trends (**Fig. S8C**,**D**). Together, these data suggested that new pole progeny may be performing more oxidative phosphorylation than older cells. To test this, we inhibited the proton motive force with the protonophore CCCP and watched the response at the single-cell level. Consistent with our hypothesis, the growth of new pole cells was affected by CCCP, while old pole cells were largely unaffected (**Fig. 4E)**. These results phenocopy LamA overexpression (*5*), further supporting the notion that LamA inhibits oxidative phosphorylation. Together, these data suggest a model whereby mycobacteria aging at the single-cell level is associated with less reliance on oxidative phosphorylation. Consequently, in an asynchronous population of mycobacterial cells, flux through central metabolism varies from cell to cell.

## Conclusion

Despite their small size, bacteria encode diverse mechanisms to spatially structure their internal biochemical processes. For example, many bacteria rely on concentration gradients along their long axis to spatially organize the macromolecular machines that perform division (*32-35*). Additionally, at division, the concentration of secondary messengers like cyclic-di-GMP, can be asymmetrically distributed, an event needed for the pathogenic lifecycle of *P. aeruginosa (36)*. Our work shows that central metabolic processes like those that produce energy can be subcellularly orchestrated in bacteria. Further work will be needed to understand the source of the connection between fluctuations in ATP levels and growth of single cells, which in mycobacteria is mediated by LamA. We speculate that the subcellular utilization of the proton motive force is an important component of this correlation. The proton motive force is a key resource for a bacterial cell – it powers the molecular machine needed to synthesize ATP and is used by numerous integral membrane proteins to transport various substrates, including molecules that make up the cell envelope, across the plasma membrane. In bacteria that grow by incorporating new envelope along their sides, complexes that use the proton motive force to grow and make ATP spatially intermingle. In contrast, we show that these two processes are spatially distinct in mycobacteria, with the proteins needed for oxidative phosphorylation found along the sides of the bacterium and certain pumps like MmpL3 and MurJ mainly localized at the poles (*37-40*) (**Fig. 5A**). Furthermore, our data suggests that LamA transiently and stochastically interacts with both pathways. Supporting this observation are the localizations of the likely kinase and phosphatase involved. PknA and PknB, essential kinases involved in cell growth and division, are predicted to phosphorylate LamA and localize to the poles (*41, 42*); in contrast, the only known protein phosphatase in mycobacteria, PstP, localizes along the side walls and to the septum (*43, 44*). These data suggest that LamA could be constantly in motion to generate asymmetry and heterogeneity in growth and division.

**Figure 5.**
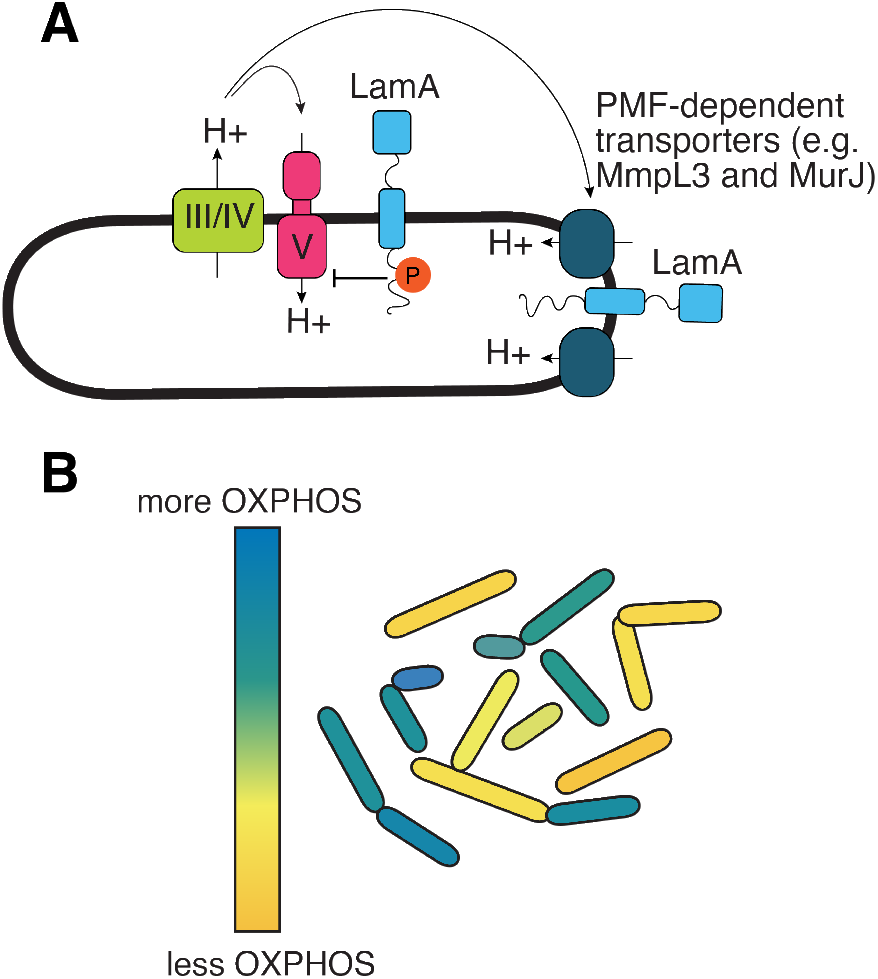
Model for spatial segregation and aging of metabolic processes in mycobacteria. **(A)** In mycobacteria, the complexes that perform oxidative phosphorylation are physically separated from those used to grow the bacterium. Both complexes rely on the proton motive force to transport molecules across the plasma membrane. LamA dynamically interacts with both complexes and is important for coupling ATP production to growth rate in single cells. **(B)** As single mycobacterial cells age, they rely less on oxidative phosphorylation for growth. Thus, in a genetically identical population of mycobacteria cells, the flux through central metabolic pathways differs from cell to cell.

Mycobacteria can simultaneously catabolize different carbon sources, a feature that is important for the survival of pathogenic mycobacteria which rely on a mixture of fatty acids, cholesterol, and carbohydrates at different points during infection (*45-47*). Our data show that the metabolism of mycobacteria is even more unusual than previously recognized, as a rapidly growing population can diversify its metabolism at the single-cell level (**Fig. 5B**). While *M. tuberculosis* is thought to mainly rely on cholesterol early in infection, multiple lines of evidence suggest that this species uses glycolysis at later stages of infection when the bacterial burden is high (*48, 49*). Our data suggest that this may also lead to metabolic and morphological heterogeneity that may be critical for the lifecycle of the pathogen. Specifically, our model suggests that inhibition of multiple metabolic subpopulations may help kill mycobacterial populations faster and more completely. Indeed, simultaneous inhibition of oxidative phosphorylation and glycolysis leads to rapid and complete sterilization of *M. tuberculosis (50)*. Thus, our data add to the growing body of evidence that understanding - and accounting for - the complexity of mycobacterial physiology at the single-cell level may be the key to improving TB therapy (*51-55*).

## Acknowledgments

We thank members of the Rego lab for thoughtful suggestions and Dr. Andy Goodman for critical reading of the manuscript.

## Funding

This work was supported by the Burroughs Wellcome Fund, the Searle Foundation, the Pew Foundation, and National Institute of Health Grant R01AI148255 to

E.H.R. We also thank Yale Flow Cytometry for their assistance with BD Symphony analyzer service. The Core is supported in part by an NCI Cancer Center Support Grant # NIH P30 CA016359. The BD Symphony was funded by shared instrument grant # NIH S10 OD026996. We also thank the MS & Proteomics Resource at Yale University for providing the necessary mass spectrometers and the accompanying biotechnology tools funded in part by the Yale School of Medicine and by the Office of The Director, National Institutes of Health (S10OD02365101A1, S10OD019967, and S10OD018034). The funders had no role in study design, data collection and analysis, decision to publish, or preparation of the manuscript.

## Author contributions

Conceptualization: C.M.G., K.R.G, and E.H.R. Investigation: C.M.G., K.R.G, and E.H.R. Resources. C.M.G., K.R.G, Y.L., L.S., and E.H.R. Software:

L.S. Supervision: E.H.R. Funding acquisition: E.H.R. Writing - original draft: C.M.G., K.R.G, and E.H.R. Writing – review and editing: C.M.G., K.R.G, Y.L., L.S., and E.H.R

## Competing interests

The authors declare no competing interests.

## Data and materials availability

Strains developed during the study will be made available from E.H.R.

## Notes

### Competing Interest Statement

The authors have declared no competing interest.

### Summary of Updates

author names corrected

## References

1. J. N. Carey et al., Regulated Stochasticity in a Bacterial Signaling Network Permits Tolerance to a Rapid Environmental Change. Cell 173, 196–207.e114 (2018).

2. N. Q. Balaban, J. Merrin, R. Chait, L. Kowalik, S. Leibler, Bacterial Persistence as a Phenotypic Switch. Science 305, 1622–1625 (2004).

3. E. S. Chung, W. C. Johnson, B. B. Aldridge, Types and functions of heterogeneity in mycobacteria. Nature Reviews Microbiology 20, 529–541 (2022).

4. B. B. Aldridge et al., Asymmetry and Aging of Mycobacterial Cells Lead to Variable Growth and Antibiotic Susceptibility. Science 335, 100–104 (2012).

5. E. H. Rego, R. E. Audette, E. J. Rubin, Deletion of a mycobacterial divisome factor collapses single-cell phenotypic heterogeneity. Nature 546, 153–157 (2017).

6. K. R. Gupta et al., An essential periplasmic protein coordinates lipid trafficking and is required for asymmetric polar growth in mycobacteria. eLife 11, e80395 (2022).

7. M. Campos et al., A constant size extension drives bacterial cell size homeostasis. Cell 159, 1433–1446 (2014).

8. J. Zeng et al., Protein kinases PknA and PknB independently and coordinately regulate essential Mycobacterium tuberculosis physiologies and antimicrobial susceptibility. PLoS Pathog 16, e1008452 (2020).

9. S. Fortuin et al., Phosphoproteomics analysis of a clinical Mycobacterium tuberculosis Beijing isolate: expanding the mycobacterial phosphoproteome catalog. Front Microbiol 6, 6 (2015).

10. K. C. Nakedi et al., Identification of Novel Physiological Substrates of Mycobacterium bovis BCG Protein Kinase G (PknG) by Label-free Quantitative Phosphoproteomics. Mol Cell Proteomics 17, 1365–1377 (2018).

11. S. Prisic et al., Extensive phosphorylation with overlapping specificity by Mycobacterium tuberculosis serine/threonine protein kinases. Proceedings of the National Academy of Sciences 107, 7521–7526 (2010).

12. R. Verma et al., Quantitative Proteomic and Phosphoproteomic Analysis of H37Ra and H37Rv Strains of Mycobacterium tuberculosis. Journal of Proteome Research 16, 1632–1645 (2017).

13. U. Kusebauch et al., <i>Mycobacterium tuberculosis</i> supports protein tyrosine phosphorylation. Proceedings of the National Academy of Sciences 111, 9265–9270 (2014).

14. T. Falk et al., U-Net: deep learning for cell counting, detection, and morphometry. Nature Methods 16, 67–70 (2019).

15. S. Berg et al., ilastik: interactive machine learning for (bio)image analysis. Nature Methods 16, 1226–1232 (2019).

16. K. C. Murphy. (Springer US, 2021), pp. 301–321.

17. D. J. Kiviet et al., Stochasticity of metabolism and growth at the single-cell level. Nature 514, 376–379 (2014).

18. M. M. Logsdon et al., A Parallel Adder Coordinates Mycobacterial Cell-Cycle Progression and Cell-Size Homeostasis in the Context of Asymmetric Growth and Organization. Current biology : CB 27, 3367–3374.e3367 (2017).

19. M. Priestman, P. Thomas, B. D. Robertson, V. Shahrezaei, Mycobacteria Modify Their Cell Size Control under Sub-Optimal Carbon Sources. Front Cell Dev Biol 5, 64 (2017).

20. E. S. Chung, P. Kar, M. Kamkaew, A. Amir, B. B. Aldridge, Mycobacterium tuberculosis grows linearly at the single-cell level with larger variability than model organisms. bioRxiv, (2023).

21. J. M. Benarroch, M. Asally, The Microbiologist’s Guide to Membrane Potential Dynamics. Trends in Microbiology 28, 304–314 (2020).

22. J. M. Kralj, D. R. Hochbaum, A. D. Douglass, A. E. Cohen, Electrical Spiking in Escherichia coli Probed with a Fluorescent Voltage-Indicating Protein. Science 333, 345–348 (2011).

23. H. Guo et al., Structure of mycobacterial ATP synthase bound to the tuberculosis drug bedaquiline. Nature 589, 143–147 (2021).

24. C. F. Wong et al., Structural Elements Involved in ATP Hydrolysis Inhibition and ATP Synthesis of Tuberculosis and Nontuberculous Mycobacterial F-ATP Synthase Decipher New Targets for Inhibitors. Antimicrobial Agents and Chemotherapy 66, e01056–01022 (2022).

25. M. G. Montgomery, J. Petri, T. E. Spikes, J. E. Walker, Structure of the ATP synthase from Mycobacterium smegmatis provides targets for treating tuberculosis. Proceedings of the National Academy of Sciences 118, e2111899118 (2021).

26. H. Yaginuma et al., Diversity in ATP concentrations in a single bacterial cell population revealed by quantitative single-cell imaging. Scientific Reports 4, 6522 (2014).

27. S. Manuse et al., Bacterial persisters are a stochastically formed subpopulation of low-energy cells. PLOS Biology 19, e3001194 (2021).

28. W.-H. Lin, C. Jacobs-Wagner, Connecting single-cell ATP dynamics to overflow metabolism, cell growth, and the cell cycle in Escherichia coli. Current Biology 32, 3911–3924.e3914 (2022).

29. K. Ginda et al., The studies of ParA and ParB dynamics reveal asymmetry of chromosome segregation in mycobacteria. Molecular Microbiology 105, 453–468 (2017).

30. M. M. Logsdon et al., A Parallel Adder Coordinates Mycobacterial Cell-Cycle Progression and Cell-Size Homeostasis in the Context of Asymmetric Growth and Organization. Current Biology 27, 3367–3374.e3367 (2017).

31. M. T. M. Hannebelle et al., A biphasic growth model for cell pole elongation in mycobacteria. Nature Communications 11, (2020).

32. Y. E. Chen et al., Spatial gradient of protein phosphorylation underlies replicative asymmetry in a bacterium. Proceedings of the National Academy of Sciences 108, 1052–1057 (2011).

33. M. Thanbichler, L. Shapiro, MipZ, a Spatial Regulator Coordinating Chromosome Segregation with Cell Division in Caulobacter. Cell 126, 147–162 (2006).

34. S. Kretschmer, P. Schwille, Pattern formation on membranes and its role in bacterial cell division. Current Opinion in Cell Biology 38, 52–59 (2016).

35. D. Kiekebusch, M. Thanbichler, Spatiotemporal organization of microbial cells by protein concentration gradients. Trends in Microbiology 22, 65–73 (2014).

36. M. Christen et al., Asymmetrical Distribution of the Second Messenger c-di-GMP upon Bacterial Cell Division. Science 328, 1295–1297 (2010).

37. C. Carel et al., Mycobacterium tuberculosis Proteins Involved in Mycolic Acid Synthesis and Transport Localize Dynamically to the Old Growing Pole and Septum. PLoS ONE 9, e97148 (2014).

38. J. Zhu et al., Spatiotemporal localization of proteins in mycobacteria. Cell Reports 37, 110154 (2021).

39. C. L. Gee et al., A Phosphorylated Pseudokinase Complex Controls Cell Wall Synthesis in Mycobacteria. Science Signaling 5, ra7–ra7 (2012).

40. L. Thouvenel, J. Rech, C. Guilhot, J.-Y. Bouet, C. Chalut, In vivo imaging of MmpL transporters reveals distinct subcellular locations for export of mycolic acids and non-essential trehalose polyphleates in the mycobacterial outer membrane. Scientific Reports 13, (2023).

41. M. Mir et al., The extracytoplasmic domain of the Mycobacterium tuberculosis Ser/Thr kinase PknB binds specific muropeptides and is required for PknB localization. PLoS Pathog 7, e1002182 (2011).

42. S. N. Nagarajan et al., Protein kinase A (PknA) of Mycobacterium tuberculosis is independently activated and is critical for growth in vitro and survival of the pathogen in the host. J Biol Chem 290, 9626–9645 (2015).

43. Iswahyudi et al., Mycobacterial phosphatase PstP regulates global serine threonine phosphorylation and cell division. Scientific Reports 9, (2019).

44. F. Shamma, E. H. Rego, C. C. Boutte, Mycobacterial serine/threonine phosphatase <scp>PstP</scp> is phosphoregulated and localized to mediate control of cell wall metabolism. Molecular Microbiology 118, 47–60 (2022).

45. A. D. Baughn, K. Y. Rhee, Metabolomics of Central Carbon Metabolism in Mycobacterium tuberculosis. Microbiology Spectrum 2, 10.1128/microbiolspec.mgm1122-0026-2013 (2014).

46. L. P. de Carvalho et al., Metabolomics of Mycobacterium tuberculosis reveals compartmentalized co-catabolism of carbon substrates. Chem Biol 17, 1122–1131 (2010).

47. Dany J. V. Beste et al., 13C-Flux Spectral Analysis of Host-Pathogen Metabolism Reveals a Mixed Diet for Intracellular Mycobacterium tuberculosis. Chemistry & Biology 20, 1012–1021 (2013).

48. J. Marrero, C. Trujillo, K. Y. Rhee, S. Ehrt, Glucose phosphorylation is required for Mycobacterium tuberculosis persistence in mice. PLoS Pathog 9, e1003116 (2013).

49. W. Y. Phong et al., Characterization of phosphofructokinase activity in Mycobacterium tuberculosis reveals that a functional glycolytic carbon flow is necessary to limit the accumulation of toxic metabolic intermediates under hypoxia. PLoS One 8, e56037 (2013).

50. J. S. Mackenzie et al., Bedaquiline reprograms central metabolism to reveal glycolytic vulnerability in Mycobacterium tuberculosis. Nat Commun 11, 6092 (2020).

51. C. Toniolo, O. Rutschmann, J. D. McKinney, Do chance encounters between heterogeneous cells shape the outcome of tuberculosis infections? Current Opinion in Microbiology 59, 72–78 (2021).

52. A. M. Cadena, S. M. Fortune, J. L. Flynn, Heterogeneity in tuberculosis. Nature reviews.Immunology, (2017).

53. P. L. Lin et al., Sterilization of granulomas is common in active and latent tuberculosis despite within-host variability in bacterial killing. Nature medicine 20, 75–79 (2014).

54. Q. Liu et al., Tuberculosis treatment failure associated with evolution of antibiotic resilience. Science 378, 1111–1118 (2022).

55. M. Mistretta, N. Gangneux, G. Manina, Microfluidic dose-response platform to track the dynamics of drug response in single mycobacterial cells. Sci Rep 12, 19578 (2022).

