## Supplemental methods and figures for "Spatial segregation and aging of metabolic processes underlie phenotypic heterogeneity in mycobacteria"

**Bacterial strains and culture conditions**. *M. smegmatis* $mc^2155$ was grown in Middlebrook 7H9 broth supplemented with 0.05% Tween80, 0.2% glycerol, 5 gm/L albumin, 2 gm/L dextrose and 0.003 gm/L catalase or plated on LB agar. For minimal medium experiments, *M. smegmatis* was grown in modified Hartmans de Bont medium (HdB) supplemented with 0.2% succinate (*1*). *M. bovis BCG* was grown in 7H9 broth supplemented with 0.05% tyloxapol, 0.2% glycerol, and 10% OADC (Fisher, #212351) or, for minimal medium experiments, MMAT broth (0.5 g/liter asparagine, 1.0 g/liter $KH_2PO_4$, 2.5 g/liter $Na_2HPO_4$, 50 mg/liter ferric ammonium citrate, 0.5 g/liter $MgSO_4·7H_2O$, 0.5 mg/liter $CaCl_2$, 0.1 mg/liter $ZnSO_4$, 10 mm glycerol, 100mM MOPS) supplemented with 0.2% succinate (*2, 3*). *Escherichia coli* DH5α cells were grown in LB broth or on LB agar plates. Concentrations of antibiotics used for *M. smegmatis* are as follows: 20 μg/ml zeocin, 25 μg/ml kanamycin, 50 μg/ml hygromycin, and 20 μg/ml nourseothricin. Concentrations of antibiotics used for *E. coli* are as follows: 40 μg/ml zeocin, 50 μg/ml kanamycin, 100 μg/ml hygromycin, and 40 μg/ml nourseothricin. All bacteria were cultured at 37˚C, with liquid cultures either shaking at 180rpm or rotating, except for the QUEEN-2m experiments, for which the bacteria were cultured at 25˚C to allow for folding of the QUEEN-2m biosensor (*4*).

**Plasmid construction and strain generation.**

*Integrating plasmids and associated strains.* All plasmids were generated via isothermal Gibson assembly whereby insert and vector backbone shared 20-25bp of homology. For analysis of LamA phospho-variants, *lamA* (MSMEG_4265) and point mutants were cloned into an L5 integrating vector under the control of a constitutive UV15 promoter, in single copy in a *lamA* deletion background. For expression of LamA controlled by the native promoter, 175bp upstream of the chromosomal locus of MSMEG_4265 was used, in line with the transcription start site dataset (*5*). For the AtpG co-IP experiment, the gene was cloned from the *M. smegmatis* genome, expressed from a constitutive UV15 promoter, and integrated as a merodiploid at the L5 site, in addition to a single copy of LamA integrated at the tweety site and driven by the UV15 promoter. For the two-color SIM experiment, the QcrB-msfGFP chromosomal fusion is described below, and Wag31-mKate2 was generated as follows. Wag31 was amplified from the *M. smegmatis* genome, expressed from the constitutive intermediate strength pTB21 promoter, and N-terminally fused to mKate2 fluorescent protein. This construct was integrated as a merodiploid at the phage Tweety site into the QcrB-msfGFP chromosomal fusion background.

*Fusions via RecET Recombineering.* To make the QcrB-msfGFP chromosomal fusion, RecET recombineering was used (*6*). To encode the fusion, a dsDNA substrate was designed using homology to the C-terminus of QcrB, an in-frame sequence coding for msfGFP, a hygromycin resistance cassette flanked by loxP sites, and homology to the region downstream of QcrB. A strain of *M. smegmatis* carrying the episomal RecET plasmid was induced overnight with isovaleronitrile and electroporated with the dsDNA substrate. Colonies were selected via hygromycin resistance and verified by PCR and microscopy.

*Oligo-mediated Recombineering.* To make protein fusions or point mutations at the native locus, we followed the protocols outlined in Murphy et al 2018 and Murphy et al 2021, respectively (*7, 8*). For making the AtpA truncation ($AtpA_{\Delta C}$), the ORBIT method was used. Briefly, a strain containing the Che9c plasmid (pKM461) was transformed with 1) a targeting oligo to the gene of interest and 2) a plasmid containing the C-terminal tag. Validation of these strains was performed by antibiotic resistance and by sequencing.

The $AtpA_{\Delta C}$ strain went through two successive rounds of recombineering, selection, and validation due to the chromosomal duplication at this locus (*9*). The targeting oligo was designed to create an insertion, and thereby truncation starting at residue V519, through the end of the protein at residue A548, following the guidance in Murphy et al 2021 (*8*). The same targeting oligo was used for both transformations, and payload plasmids pKM491 ($hyg^R$) and pKM496 ($zeo^R$) were used in successive rounds of ORBIT.

For creating SNPs at the native locus, we used a strain containing a mutated hygromycin resistance cassette integrated at L5 as the base (pKM427). This strain was transformed with 1) a targeting oligo containing the mutation and 2) an oligo containing an SNP that repairs the mutated hygromycin cassette. Validation of these strains was performed by $Hyg^R$ screening and by sequencing. The $LamA_{Y50A}$ strain was generated with this protocol.

**Conventional Microscopy.** An inverted Nikon Ti-E microscope was used for the time-lapse and snapshot imaging. An environmental chamber (Okolabs) maintained the samples at 37˚C, except QUEEN experiments which were maintained at 25˚C. To reduce phototoxicity, exposure times were kept below 100ms for excitation with 470nm and 300ms for excitation with 550nm.

*Timelapse*. Exponentially growing cells were cultivated in a B04 microfluidics plate from CellAsic (Millipore), continuously supplied with fresh 7H9 or minimal medium, where described, and imaged every 15 minutes using a 60x 1.4 N.A. Plan Apochromat phase contrast objective (Nikon). For BCG timelapse, the imaging setup was the same except phase images were captured every 1 hour for 5 days. Fluorescence was excited using the Spectra X Light Engine (Lumencor), separated using single- or multi- wavelength dichroic mirrors, filtered through single bandpass emission filters, and detected with an sCMOS camera (ORCA Flash 4.0). Filters are: GFP (Ex: 470/24; Em: 515/30); QUEEN (Ex: 395/25; Em: 515/30); mKate2 (Ex: 550/15; 595/30). To reduce phototoxicity exposure times were kept below 100ms for excitation with 395nm and 470nm, and 300ms for excitation with 550nm. For CCCP growth inhibition timelapse (Fig 4E), 7H9 media was continuously supplied to the imaging chamber for either 4 or 10 hours, before switching to 7H9 supplemented with 50µM CCCP for the duration of the timelapse.

*Snapshot and agar pad imaging.* Exponentially growing cells were prepared as described above. Agar pads containing 1% UltraPure Agarose (ThermoFisher, #16500500) were made with

HdB base media lacking tyloxapol and carbon sources. Cells were immobilized under pads, and a humidifier was used for short time-lapses.

**Image analysis.**

*U-Net segmentation*. Phase contrast and fluorescence time-lapse images were analyzed in open-source image analysis software Fiji (*10*). Images were processed with a custom pipeline using Fiji, iLastik (*11*), and U-Net (*12*). The U-Net plugin in FIJI was used as the platform for training and segmenting our phase-contrast microscopy images. Instead of using the raw images as the input, however, we first obtained the Hessian of Gaussian Eigenvalues (HoGE, with σ value of 0.7) of the raw images, either with the iLastik software or a custom MATLAB script translated from the relevant iLastik source code. Using the HoGE images as the input, we trained our U-Net model using about ten 2D images and their associated segmentation masks manually generated and the trained model was subsequently used in all our U-Net segmentation tasks.

Output segmented files were then transferred to Fiji where we used the binary masks to manually annotate cells using the magic wand tool. Basic analysis (Figs 1C/D, 4E, S1, S2) was done with direct measurement of birth length, width, area, and mean fluorescent signal in Fiji. Further analysis was done with custom MATLAB scripts for generating: exponential elongation rate measurements (Figs 1E, 3C, S3B), fluorescent line profiles (Fig 1H, S4, S5B); and custom R scripts for generating: division asymmetry (Figs 1D for asymmetry of area; 4B for asymmetry of fluorescent signal) and aging analyses (Figs 4C, S8).

*QUEEN-2m analysis*. To analyze the QUEEN-2m data shown in Fig 3, we collected fluorescence using two filter configurations. GFP (Ex: 470/24; Em: 515/30); QUEEN (Ex: 395/25; Em: 515/30). The resulting time-lapse stacks were background subtracted, and individual and pixel-by-pixel “ratio images” were generated by division of the image collected by the QUEEN filter divided by the image collected by the GFP filter. Associated phase contrast images were segmented using the U-Net pipeline described above, and these masks were multiplied with the “ratio image”. Next, individual cell cycles were tracked from birth to division, and both the cell area and QUEEN-2m signal were recorded at each time point. For each cell cycle, the following parameters were measured: (1) the exponential growth rate, $\lambda$, was extracted by fitting the cell-area curve to an exponential, and (2) ATP variation signal was calculated by normalizing the standard deviation of the QUEEN-2m signal to the average signal per cell cycle.

*Aging analysis.* Cells were initially chosen at a division event that resulted in a 0-1 pole age cell (see Fig 4A). This was taken as the mother cell and annotated as time = 0. Successive rounds of division were followed, following the age=1 pole of that initial mother cell. This was done with a custom lineage tracing MATLAB script which takes the mask of the birth cell chosen with the magic wand tool as input and follows until it finds a division event. The output of this is individual ROIs that correspond to cell masks at each time point from birth to division. These ROIs are then taken to the corresponding fluorescent image and measured for mean fluorescence intensity and time slice.

To generate an average profile of multiple cells, both time since birth and mean fluorescence intensity were normalized using a custom R script. Briefly, this was done by slicing the total division time equally, controlling for total generation time to generate the “pole age” metric, and by setting the fluorescence at t=0 as the maximum. To generate Fig S8A/C, fluorescence values for all cells at t=0,1,2,3 were plotted. To generate Fig S8B/D, cells displaying the “step down” phenotype – defined as cells in which fluorescence at t=final was less than at t=0 – were pooled and a LOWESS regression with 20-point “fine” smoothing was performed independently for each biological replicate using Prism 10 and plotted.

Similarly, for Fig 4D cells were chosen based on lineages that could be tracked for three divisions. The initial mother cell was chosen agnostic to its initial pole age. These were subsequently followed to quantify mean QcrB-msfGFP fluorescence intensity at birth. Ten separate lineages were measured, and cells were matched for their pole age identity at birth.

These matched cells were averaged, and each of these values was normalized to the maximum fluorescence intensity of the entire average lineage.

*BCG analysis*. M. bovis BCG cells were morphologically different enough from M. smegmatis, especially in the minimal media condition, that we could not use our U-Net training set to segment these images. Instead, the HoGE images, as described above, were used to manually measure cell length.

*Single-cell response to CCCP.* Images were pre-processed with the U-Net pipeline described above. Daughter cells were chosen and annotated for old or new pole identity, and their areas measured directly in Fiji. Cells which divided during the 7H9 only "-CCCP" condition were counted only if the division event occurred before the media was switched. For the "+CCCP" condition, divisions were only counted if the event occurred at or greater than 30 minutes after the beginning of flowing CCCP supplemented media. This rule was chosen to allow equal circulation of drug into the imaging chamber.

**Three-dimensional structured illumination microscopy.** 3D-SIM imaging shown in Figure 2 was performed on our custom live-cell 3D structured-illumination microscope based on previously published methods (*13, 14*). One major modification pertains to how the *s*-polarization of the illumination beams, essential for creating high-contrast SIM pattern and thus high-quality images, was maintained for all SIM angles. After a spatial-light modulator diffracts the incoming laser beam, the linearly polarized beams were first passed through a quarter-wave plate and became circularly polarized, and then were directed through a custom-made round-shaped compound linear polarizer (Laser Components) composed of 12 equally spaced sectors and a small central circular area. While the central area is non-polarizing, each sector linearly polarizes along the direction tangential to the midpoint of its arc (*15*). Hence for each SIM orientation, the side beams are both linearly polarized perpendicular to the plane of incidence (i.e., *s*-polarized) and the central beam remained circularly polarized. Even though the central beam is not colinearly polarized with the side beams, we found that in practice this method is sufficient in creating high-contrast first-order 3D SIM illumination patterns. The main benefit is the zero time delay as opposed to the 10s of milliseconds delay typical for liquid crystal polarization rotating devices.

**AtpG-LamA coIP.** Mycobacterial cultures were grown to mid-log phase as described above. Protocol for crosslinking was adapted from Belardinelli et al, Sci Rep, 2019 (*16*). Cells were washed with 1x phosphate buffered saline (PBS) once. Pellets were resuspended in 1mL of 1x PBS with 1.25mM dithiobis (succinimidyl propionate) (DSP) and incubated for 30 minutes rolling at 37˚C for crosslinking. After incubation, cells were pelleted at 10,000xg for 5 minutes at room temperature and the supernatant was discarded. The pellet was resuspended in lysis buffer (50mM Tris-HCl, pH 7.4; 150mM NaCl; 10ug/mL DNase I; one tablet Roche cOmplete EDTA free protease inhibitor cocktail; and and 0.5% Igepal Nonidet P40 Substitute) and lysed with a Bullet Blender Homogenizer (Next Advance) on speed 8 for 5 minutes thrice in a cold room. Lysed cells were spun down at 15,000xg for 15 minutes at 4˚C, and the supernatant was transferred to a clean eppendorf tube. Lysates were incubated, where indicated, with either magnetic a-FLAG M2 beads (Sigma-Aldrich, #M8823) or magnetic Pierce Streptavidin beads (Thermo Scientific, #88816) and incubated at 4˚C overnight, rotating. After incubation, samples were spun down at 2,500xg for 1 minute at room temperature and flow through was discarded. Beads were washed three times with non-detergent wash buffer (10mM Tris-HCl, pH 7.4; 150mM NaCl, 0.5mM EDTA). FLAG M2 beads were eluted twice with 3xFLAG peptide (Sigma-Aldrich, #F4799) for 30 minutes rotating at 4˚C. Streptavidin beads were eluted with 1x BXT buffer (IBA) for 10 minutes at room temperature. All samples were prepared for western blotting by addition of Laemmli Buffer + DTT and boiled at 95˚C for 5 minutes, to reverse all crosslinks.

*Western blot verification.* Samples were run on NuPAGE™ (Thermo Fisher) 4-12% Bis-Tris gels in 1x MOPS-SDS running buffer. Proteins were transferred to nitrocellulose membranes and probed with 1:1,000 $\alpha$-FLAG primary (Sigma-Aldrich, clone M2) and 1:5,000 $\alpha$-mouse secondary (Thermo Fisher, Superclonal™ A28177). SuperSignal™ West Femto Extended Duration Substrate (Thermo Fisher, #34094) was used for chemiluminescent visualization on an Amersham ImageQuant 800 system.

**Mass spectrometry.** Precipitation of LamA-3xFLAG was prepared as described above, with the following modifications. Crosslinking was done with 1% PFA in PBST for 1 hour, rolling at 37˚C. Cells were lysed in lysis buffer (50mM Tris-HCl pH 7.4, 50mM NaCl, 1% DDM, and one tablet Roche cOmplete EDTA free protease inhibitor cocktail) and lysed with a BeadBug™ Microtube Homogenizer at 4,000rpm six times for 30 seconds each, icing in between. Wash buffer was prepared as follows: 100mM Tris, pH 8.0, 150mM NaCl, 1mM EDTA. After verification by western blot, performed as above, samples were sent to the Yale Keck MS & Proteomics Core for in-gel digestion with Trypsin and GluC, and analyzed with a Thermo Scientific LTQ-Orbitrap XL mass spectrometer. Spectra were determined in-house using the Mascot algorithm and Mascot Distiller program.

**Flow cytometry**. Mycobacterial cultures grown to mid-log phase were diluted to an OD = 0.3. As appropriate, cells were first incubated with 500µM CCCP for 15 minutes rolling at 37˚C. Cells were then stained with 0.2µM Image-iT™ TMRM (Thermo Fisher, #I34361) for 30 minutes rolling at 37˚C. Untreated and unstained controls were incubated alongside to control for growth. All samples were analyzed on a BD Symphony Flow Cytometer with channels as follows: FITC (488nm laser, 200mW, 515/25nm bandpass, 505nm long pass filters) and PE (561nm laser, 150mW, 586/15nm bandpass filter). Data was analyzed using FlowJo™ 10.6.2 Software (BD Life Sciences). Doublet discrimination was performed with cultures of GFP-expressing, mScarlet-expressing, or overnight mixed co-cultures of mycobacteria, where a threshold of <2% doublets was applied. This gating strategy was then applied to the experimental samples uniformly to capture only single cells.

**Bulk ATP measurement.** ATP levels in whole cells were measured using luminescence based BacTiter-Glo™ microbial cell viability kit (Promega, #G8231). The kit contains a thermostable luciferase which reacts with the ATP molecules present inside the cell to produce luminescence. Briefly, 100µL log phase mycobacterial culture was mixed with 100µL BacTiter-Glo reagent in a 96 well flat black Nunc™ plate (Thermo Scientific, #12-566-72) and incubated at room temperature for 30 minutes. Luminescence readings were acquired in a Tecan Spark® multi-mode microplate reader. Normalization of readings was done by dividing the luminescence values with the $OD_{600}$ value of the bulk bacterial culture. All assays were performed in biological triplicates.

**Experimental replicates.** Biological replicates are defined as independent cultures grown in parallel or on separate days. Technical replicates are defined as the same culture, measured independently. All the experiments were performed at least twice, with biological replicates, except for the *M. bovis* BCG experiment, which was performed once.

**Table S1. Possible interaction partners.** Unique proteins that pull down with LamA as bait. Hits fall into two large categories – cell growth and division, and oxidative phosphorylation.

| Protein Name | Gene ID | Function |
|---|---|---|
| PonA1 | MSMEG_6900 | Essential bi-functional peptidoglycan synthase |
| MurA | MSMEG_4932 | Essential lipid II precursor synthesis |
| UmaA | MSMEG_0913 | Methoxy mycolic acid synthesis |
| PgfA | MSMEG_0317 | Essential protein involved in mycolic acid biosynthesis |
| MmpL3 | MSMEG_0250 | Essential protein involved in mycolic acid biosynthesis |
| PknA | MSMEG_0030 | Essential kinase involved in growth and division |
| N/A | MSMEG_3058 | Unknown lipoprotein |
| QcrA | MSMEG_4262 | Fe-S subunit of cytochrome bc1 complex |
| QcrB | MSMEG_4263 | Cytochrome bc1 complex subunit b |
| CtaC | MSMEG_4268 | Cytochrome aa3 subunit 2 |
| AtpA | MSMEG_4938 | Alpha subunit of ATP synthase |
| AtpB | MSMEG_4942 | A subunit of ATP synthase |
| AtpFH | MSMEG_4939 | Beta-delta subunit of ATP synthase |
| AtpG | MSMEG_4937 | Gamma subunit of ATP synthase |

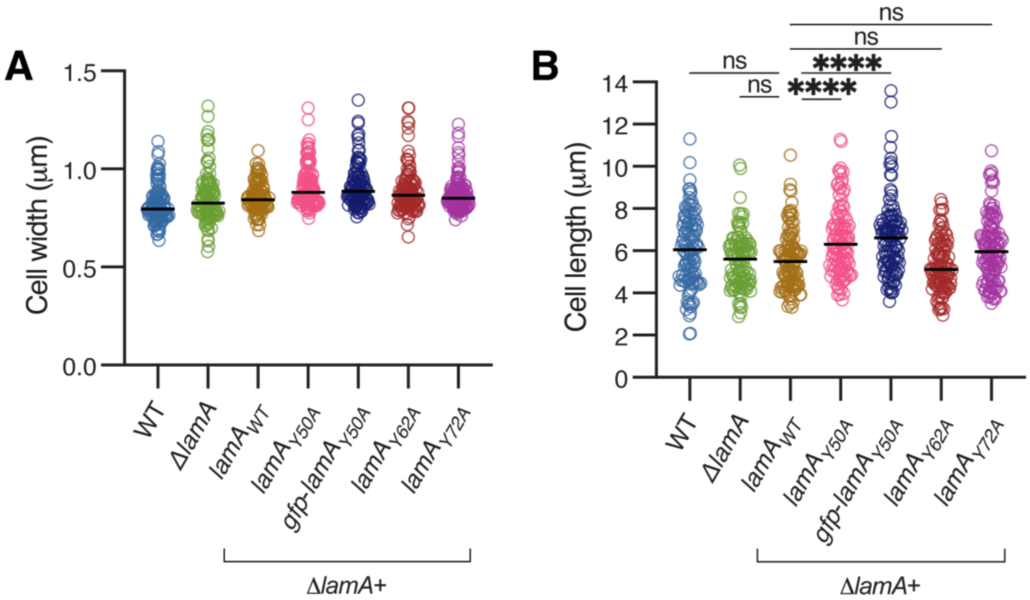


**Figure S1. Length and width measurements of LamA phosphovariants.** Cells were segmented with the custom U-net pipeline, and various metrics were measured using Fiji. Dark black lines indicate medians. **(A)** Cell widths of WT, Δ*lamA*, or cells expressing the indicated *lamA* allele in Δ*lamA*; n=100-125 cells for all strains. **(B)** Cell lengths of WT, Δ*lamA*, or cells expressing the indicated *lamA* allele in Δ*lamA*; n=100-125 cells for all strains; **** p<0.0001 by one-way ANOVA comparing the means of all strains to the Δ*lamA* + $lamA_{WT}$ strain.

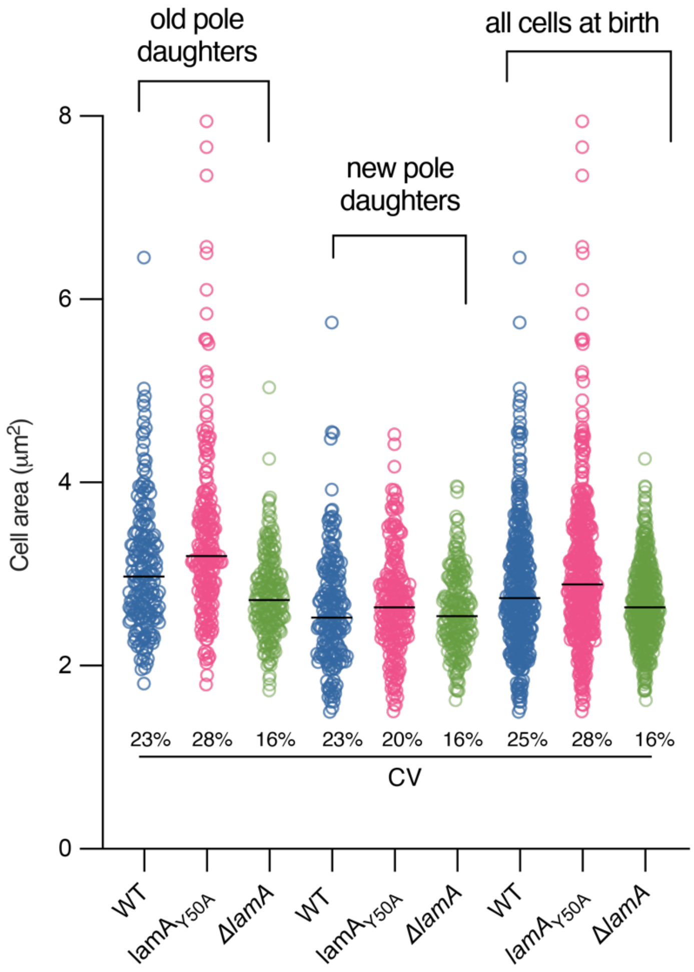


**Figure S2. A conserved phosphorylated tyrosine of LamA is important for variability in mycobacteria.** Cell areas were measured in WT, Δ*lamA*, and the chromosomally mutated *lamA*$_{Y50A}$ for cells at birth, and grouped by pole identity from the previous division. Black lines indicate medians; CV = coefficient of variation; n = 195-235 cells for all strains.

**A**

| Metric | equation | CV for WT | CV for *ΔlamA* | CV for $LamA_{Y50A}$ |
|---|---|---|---|---|
| **Exponential growth rate** | $\lambda = \ln(S_d/S_b)/\Delta T$ | 16% | 11% | 10% |
| **Size normalized growth rate** | $(S_d - S_b)/(S_b * \Delta T)$ | 25% | 16% | 17% |
| **Linear growth rate** | $(S_d - S_b)/\Delta T$ | 23% | 16% | 18% |

$S_d$ = the size of the cell at division
$S_b$ = the size of the cell at birth
$\Delta T$ = the time between birth and division

**B**

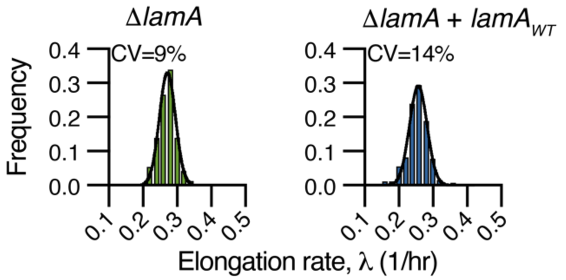


**Figure S3. Exponential growth is more homogenous in Δ*lamA* populations. (A)** Data shown in Fig. 1E are re-analyzed to compute different metrics of growth rate, and their corresponding coefficient of variation (CV) for WT, Δ*lamA*, and the chromosomal $LamA_{Y50A}$ mutant, respectively. **(B)** Exponential growth rate λ, of the indicated strains was calculated from time-lapse phase contrast microscopy videos; n = 165 and 255 cells for Δ*lamA* and Δ*lamA* + *lamA*$_{WT}$, respectively.

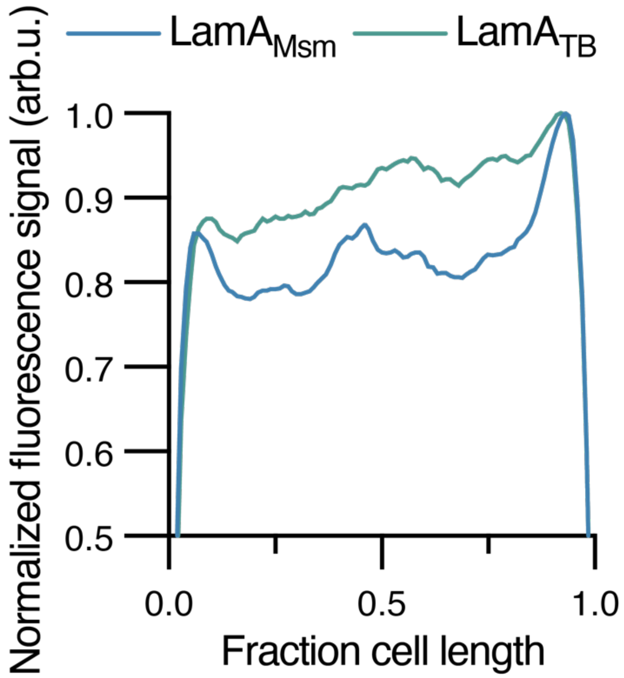


**Figure S4. LamA from *M. tuberculosis* is localized evenly across the length of the cell body.** Line profiles from phase contrast snapshot images were generated from strains containing either msfGFP-LamA$_{Msm}$ or msfGFP-LamA$_{TB}$, both expressed from a medium-strength constitutive promoter. Intensity profiles were measured across several cells, normalized by both cell length and maximum fluorescence in each cell, and averaged (n = 165 cells and 140 cells for *Msm* and *Mtb* versions, respectively).

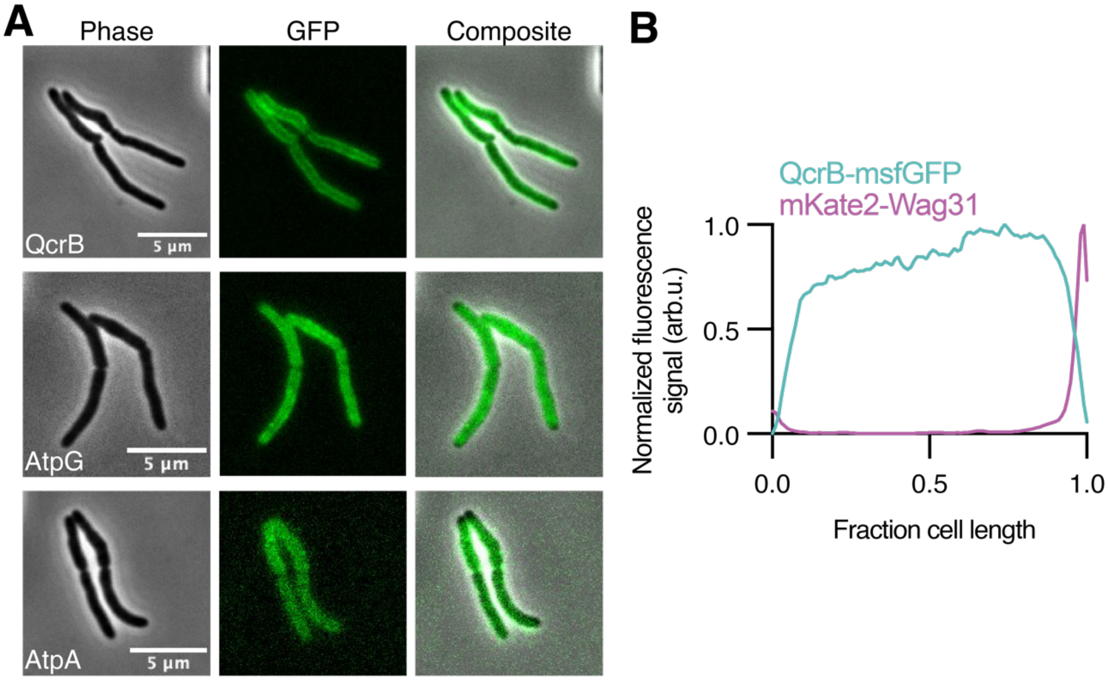


**Figure S5. Components of the electron transport chain and ATP synthase localize to the membrane. (A)** Phase contrast and fluorescence images of QcrB-msfGFP, AtpG-msfGFP, and AtpA-msfGFP. Representative images were chosen from snapshots originating from live-cell time-lapse videos of cells in CellAsic microfluidic devices. **(B)** QcrB and Wag31 are spatially segregated in individual cells. Line profiles were generated in Fiji from 3D-SIM images and aligned to the brightest pole using a custom MATLAB script; n=93 cells.

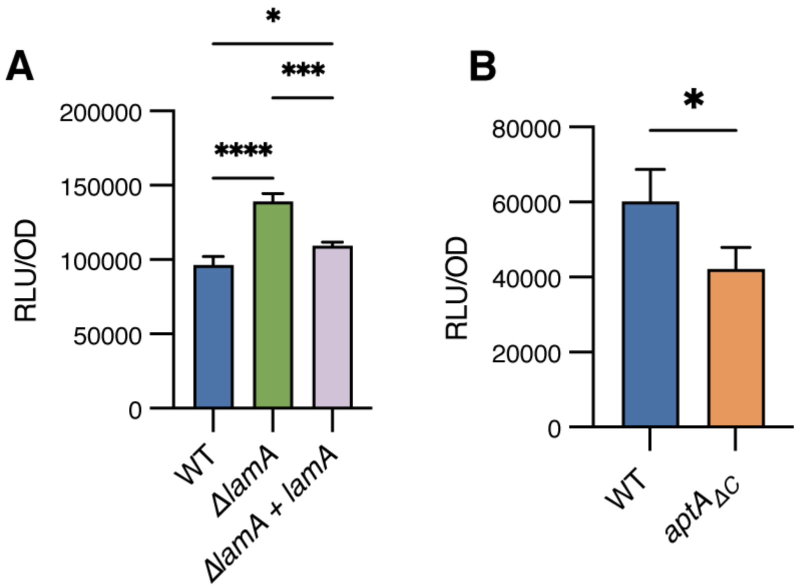


**Figure S6. ATP measurements**. **(A)** ATP levels were quantified using the BacTiter-Glo assay in WT, Δ*lamA*, and pNative *lamA* complement cells. Means are representative of biological triplicates, bars represent means, error bars represent standard deviation of the mean. *p<0.05 to show partial complementation of *lamA* driven by the native promoter; ***p<0.0005, ****p<0.0001, as tested by a one-way ANOVA with Tukey's multiple comparisons. **(B)** ATP levels were quantified using the BacTiter-Glo assay in WT and $atpA_{\Delta C}$ cells. Means are representative of biological triplicates, bars represent means, error bars represent standard deviation of the mean. *p<0.05 tested with a two-tailed students t-test.

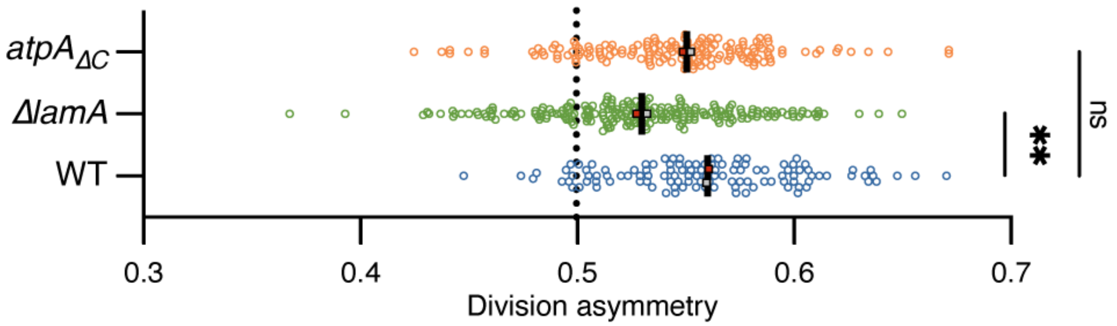


**Figure S7. Deletion of an actinobacterial-specific extension on the ATP synthase does not affect division asymmetry.** Asymmetry in the daughter cell size at the time of division, measured by the old pole progeny as a fraction of the total size of both siblings. The dotted line marks symmetric division. (n= 115, 225, and 154 sibling cell pairs for WT, Δ*lamA*, and $atpA_{\Delta C}$, from two biological replicates, respectively). Magenta and grey squares represent the medians of individual biological replicates, and the dark black line represents the median of those medians. *p<0.05, nd = no discovery, calculated by a paired ANOVA comparing to means to WT, corrected for multiple comparisons.

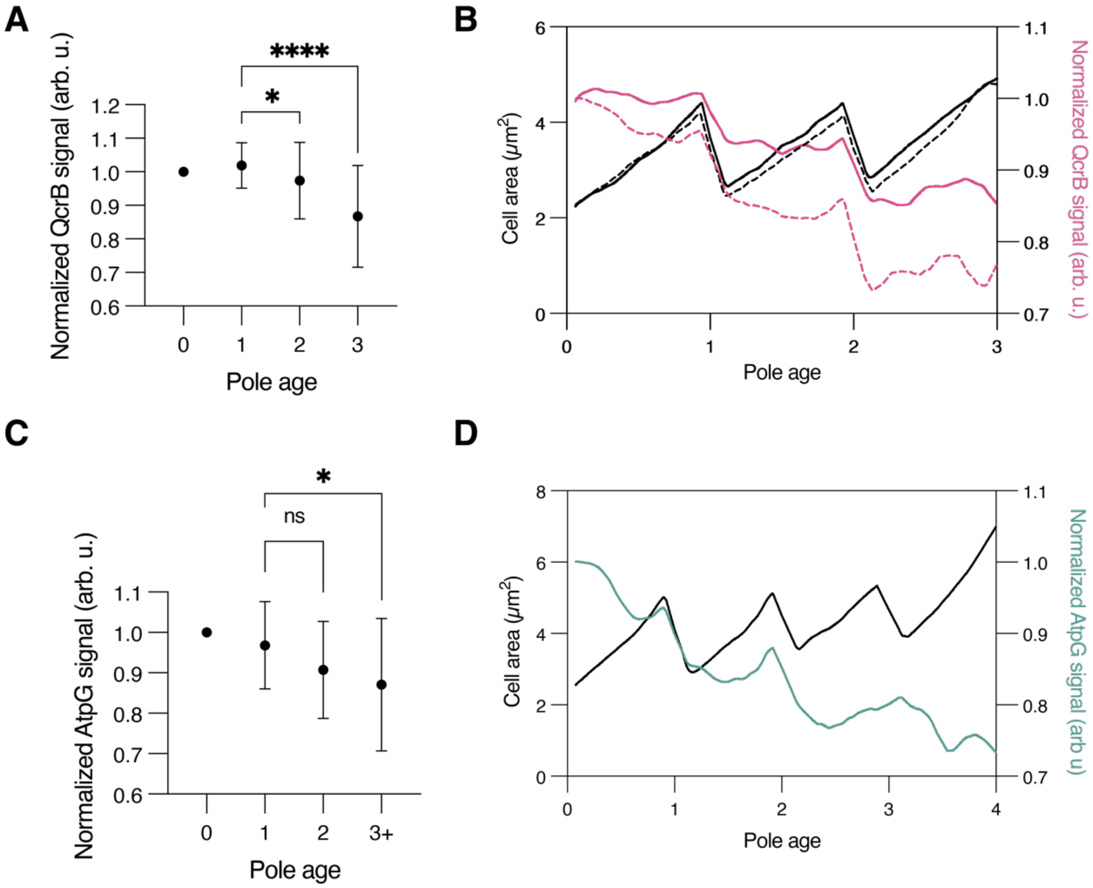


**Figure S8. QcrB and AtpG signals decrease at division events in old pole cells. (A)** (n=51 lineages from two biological replicates.) New pole cells were identified and followed for two or three divisions, tracking the oldest pole of those lineages. Pole age of zero was defined as the birth of the new pole. Fluorescent signal was normalized to the intensity at pole age zero. Dots represent mean and bars represent the standard deviation. $^*p<0.05$, $^{****}p<0.0001$, calculated by a paired ANOVA comparing the means to pole age 1, corrected for multiple comparisons. **(B)** For cells that displayed the step-down phenotype, line graphs of fluorescent signal (right y-axis) and cell size (left y-axis) were normalized to their values at pole age zero, and several lineages were averaged. Lines were generated by LOWESS regression in Prism. Solid versus dashed lines represent individual biological replicates with n= 24 and 8 lineages, respectively. **(C)** As in panel A, old pole cells expressing AtpG-msfGFP were followed over generations 2-4 generations. $^*p<0.05$, calculated by a paired ANOVA comparing the means to pole age 1, corrected for multiple comparisons (n=22 lineages from one biological replicate.) **(D)** As in panel B, old pole cells expressing AtpG-msfGFP cells were tracked over 3-4 generations, normalized, and averaged (n=14 lineages).

**Supplementary Table 1. List of strains used in this work.**

| Reagent type (species) or resource | Designation | Source or reference | Identifiers | Additional Information |
|---|---|---|---|---|
| Strain (*Mycobacterium smegmatis)* | | | *Mycobacterium smegmatis* mc$^{2}$155 | Wild type *M. smegmatis* |
| Strain (*Mycobacterium bovis*) | | Gift from John MacMicking | *Mycobacterium bovis BCG* | Wild type *M. bovis BCG,* Phipps strain, ATCC 35744 |
| Strain (*M. smegmatis*) | HR342 | Rego et al 2016 | mc$^{2}$155 ΔMSMEG_4265 | LamA knockout strain |
| Strain (*M. smegmatis*) | HR362 | Rego et al 2016 | mc$^{2}$155 ΔMSMEG_4265 L5: pNative MSMEG_4265 | WT *M. smegmatis* LamA for complementation driven by the native promoter |
| Strain (*M. smegmatis*) | HR486 | This work | mc$^{2}$155 ΔMSMEG_4265 tw::pTetO-lamA$_{WT}$-strep | WT *M. smegmatis* LamA for complementation |
| Strain (*M. smegmatis*) | HR528 | This work | mc$^{2}$155 ΔMSMEG_4265 tw:: pTetO-lamA$_{Y50A}$-strep | Integrated single copy of LamA$_{Y50A}$ in a Δ*lamA* background |
| Strain (*M. smegmatis*) | HR521 | This work | mc$^{2}$155 ΔMSMEG_4265 tw:: pTetO-GFP-lamA$_{Y50A}$-strep | Integrated single copy of GFP-LamA$_{Y50A}$ in a Δ*lamA* background |

| | | | | |
|---|---|---|---|---|
| Strain (*M. smegmatis*) | HR482 | This work | $mc^2155$ ΔMSMEG_4265 tw:: pTetO-lam$A_{Y63A}$-strep | Integrated single copy of Lam$A_{Y63A}$ in a Δ*lamA* background |
| Strain (*M. smegmatis*) | HR483 | This work | $mc^2155$ ΔMSMEG_4265 tw:: pTetO-lam$A_{Y73A}$-strep | Integrated single copy of Lam$A_{Y73A}$ in a Δ*lamA* background |
| Strain (*M. smegmatis*) | CMG590 | This work | $mc^2155$ *lam*$A_{Y50A}$ L5::Hyg-repaired-$Zeo^R$ | Oligo recombineered SNP strain with Y50A at the native locus of LamA |
| Strain (*M. smegmatis*) | CMG766 | This work | $mc^2155$ ΔMSMEG_4265 L5:: pNative-msfGFP lam$A_{WT}$ | Localization of *M. smegmatis* Lam$A_{WT}$ from the native promoter |
| Strain (*M. smegmatis*) | CMG768 | This work | $mc^2155$ ΔMSMEG_4265 L5:: pNative-msfGFP lam$A_{Y50A}$ | Localization of *M. smegmatis* Lam$A_{Y50A}$ from the native promoter |
| Strain (*M. smegmatis*) | CMG754 | This work | mc2155 ΔMSMEG_4265 L5:: ptb21-msfGFP lam$A_{Msm\ WT}$ | Localization of *M. smegmatis* LamA |
| Strain (*M. smegmatis*) | CMG750 | This work | mc2155 ΔMSMEG_4265 L5:: ptb21-msfGFP lam$A_{TB\ WT}$ | Localization of *M. tuberculosis* LamA |
| Strain (*M. smegmatis*) | CMG495 | This work | $mc^2155$ ΔMSMEG_4265 L5:: pTetO AtpG-3xFLAG ptw:: pTetO lamA-strep | Strain for coIP of AtpG with LamA |
| Strain (*M. smegmatis*) | CMG536 | This work | $mc^2155$ qcrB-msfGFP | Chromosomally tagged QcrB created with RecET recombineering |

| | | | | |
|---|---|---|---|---|
| Strain (*M. smegmatis*) | CMG713 | This work | $mc^2155$ ΔMSMEG_4265 qcrB-msfGFP | Tagged QcrB fusion in Δ*lamA* background |
| Strain (*M. smegmatis*) | CMG775 | This work | $mc^2155$ ΔMSMEG_4265 qcrB-msfGFP L5:: pTetO-$lamA_{WT}$-3x FLAG | Tagged QcrB fusion in Δ*lamA* background with $LamA_{WT}$ complement |
| Strain (*M. smegmatis*) | HR312 | This work | $mc^2155$ L5::pTetO-atpG-msfGFP | Integrated tagged AtpG |
| Strain (*M. smegmatis*) | HR311 | This work | $mc^2155$ L5::pTetO-atpA-msfGFP | Integrated tagged AtpA |
| Strain (*M. smegmatis*) | CMG809 | This work | $mc^2155$ qcrB-msfGFP tw:: ptb21 mKate2-wag31 | Two-color strain with tagged QcrB and Wag31 |
| Strain (*M. smegmatis*) | CMG737 | This work | $mc^2155$ AtpA::4xGly-TEV-Flag-6xHis AtpA-Cterm::zeoR | AtpA C-terminal truncation strain; both copies truncated; aka $AtpA_{\Delta C}$ |
| Strain (*M. smegmatis)* | CMG551 | This work | $mc^2155$ tw:: pTetO-mScarletI-FLAG | Control strain for flow cytometry |
| Strain (*M. smegmatis)* | CMG552 | This work | mc2155 giles::pTetO-msfGFP-myc | Control strain for flow cytometry |
| Strain (*M. smegmatis)* | CMG804 | This work | $mc^2155$ / pTetO-mycoQUEEN-2m | Codon optimized for *M. tuberculosis* with IDT |

| | | | | |
|---|---|---|---|---|
| Strain (*M. smegmatis)* | CMG808 | This work | $mc^2$155 ΔMSMEG_4265 / pTetO-mycoQUEEN-2m | Codon optimized for *M. tuberculosis* with IDT |
| Strain (*M. smegmatis*) | HRCT146 | This work | $mc^2$155 L5::HygSTOP-$Zeo^R$, pTet-RecT-Bxb1Int-SacB-kan | Base strain in *M. smegmatis* for oligo mediated SNP recombineering |
| Strain (*M. smegmatis*) | HRCT078 | This work | $mc^2$155 / pKM461 | Base strain in *M. smegmatis* for ORBIT |
| Plasmid (*E. coli*) | KG3 | This work | DH5α / pL5-pTetO-mycoQueen_2m | Codon optimized for *M. tuberculosis* with IDT |
| Plasmid (*E. coli*) | pKM402 | Murphy et al 2018 | DH5α / pL5 Che9c RecT $Hyg^{mutated}$ | Integrating plasmid for broken Hyg cassette |
| Plasmid (*E. coli*) | pKM496 | Murphy et al 2018 | DH5α / attB $cam^R$ oriE pEM7 $zeo^R$ | Zeo knockout plasmid for ORBIT |
| Plasmid (*E. coli*) | pKM491 | Murphy et al 2018 | DH5α / pHyg $Hyg^R$ $cam^R$ oriE attB C-terminal tag: 4xGly-TEV-Flag-6xHis | C-terminal tagged plasmid for ORBIT |
| Plasmid (*E. coli*) | CMG740 | This work | Dh5α / pL5 ptb21 msfGFP-$lamA_{TB\ WT}$ | Integrative plasmid for tagged WT LamA from *M. tuberculosis* |
| Plasmid (*E. coli*) | CMG747 | This work | Dh5α / pL5 ptb21 msfGFP-$lamA_{Msm\ WT}$ | Integrative plasmid for tagged WT LamA from *M. smegmatis* |

| | | | | |
|---|---|---|---|---|
| Plasmid (*E. coli*) | CMG719 | This work | Dh5α / pL5 pNative msfGFP-lam$A_{Msm\ WT}$ | Integrative plasmid with native promoter and tagged WT LamA from *M. smegmatis* |
| Plasmid (*E. coli*) | CMG720 | This work | Dh5α / pL5 pNative msfGFP-lam$A_{Msm\ Y50A}$ | Integrative plasmid with native promoter and tagged Lam$A_{Y50A}$ from *M. smegmatis* |
| Plasmid (*E. coli*) | CMG483 | This work | DH5α / pL5 ptetO AtpG-3xFLAG | Plasmid with 3xFLAG tagged AtpG used for coIP |
| Plasmid (*E. coli*) | CMG817 | This work | DH5α / ptw ptb21 mKate2-wag31 | Plasmid with mKate2-wag31 for SIM imaging |

**Supplementary Table 2. List of oligos used in this work.**

| Oligo Name | Sequence | Use | Type |
|---|---|---|---|
| HR408 | gcttaattaagaaggagatatacatatgagcgggccgaatcccacgg | FW primer for pTetO LamA$_{WT}$ | Primer |
| HR424 | ccccaattaattagctaaagctttcacttctcgaactgggggtggctccagtcgcag gcagtctgcgccgagttgttgg | RV primer for LamA$_{WT}$-strep | Primer |
| HR494 | ccggcgcggagcacgcctgggag | FW primer for L5 LamA$_{Y50A}$ | Primer |
| HR495 | ctcccaggcgtgctccgcgccgg | RV primer for L5 LamA$_{Y50A}$ | Primer |
| HR498 | gtcggcaggcacggcgggaccagcggtg | FW primer for L5 LamA$_{Y63A}$ | Primer |
| HR499 | caccgctggtcccgccgtgcctgccgac | RV primer for L5 LamA$_{Y63A}$ | Primer |
| HR500 | cgggtcgtagttgtcggcgtcatagagcgccacg | FW primer for L5 LamA$_{Y73A}$ | Primer |
| HR501 | cgtggcgctctatgacgccgacaactacgacccg | RV primer for L5 LamA$_{Y73A}$ | Primer |
| CMG374 | agcgggccgaatcccccg | FW primer for LamA$_{TB}$ | Primer |
| CMG371 | ggtccccaattaattagctaaagctttcagcagctcgtttgcggtc | RV primer for msfGFP-LamA$_{TB}$ Gibson, LamA piece | Primer |
| CMG372 | cgggggattcggcccgcttccagaacctttgtagag | RV primer for msfGFP-LamA$_{TB}$ Gibson, msfGFP piece | Primer |
| CMG382 | gcgtttaatactgcatgcactctagaccctcatcaccggtacgttc | FW primer for pNative lamA$_{Msm}$ | Primer |
| CMG383 | cggtgaacagttcttcacctttcatctacgcctatccttctcgac | RV primer for pNative lamA$_{Msm}$ | Primer |
| CMG384 | aactctacaaaggttctggaagcgggccgaatcccacg | FW primer for msfGFP-lamA$_{Msm,}$ lamA piece | Primer |
| CMG385 | cgtgggattcggcccgcttccagaacctttgtagagtt | RV primer for msfGFP-lamA$_{Msm}$, lamA piece | Primer |

| | | | |
|---|---|---|---|
| CMG386 | catcgatagaaaggaggaaggaaagcttatgaaaggtgaagaactgttcaccg | RV primer for msfGFP-lamA$_{Msm}$, msfGFP piece | Primer |
| CMG280 | cttaattaaggaggagatatacatatggcagccacactgcgcg | FW primer for AtpG-3xFLAG | Primer |
| CMG281 | ccgtcatggtctttgtagtcgatatctttcgagccggccagcgc | RV primer for AtpG-3xFLAG | Primer |
| CMG255 | cctctagggtccccaattaattagctaaagctttcacttatcatcgtcatctttatag | RV primer for 3xFLAG | Primer |
| CMG126 | gcgtgcggccgctgattagctaagctaatattggatcgtcgcacc | FW primer for ptb21 | Primer |
| CMG132 | cagttcttcacctttcataagctttccttcctcctttctatcgatggatccgtggcaaga gggtc | RV primer for ptb21-msfGFP | Primer |
| CMG428 | ccttaatcagctcagacaccatatgttaattaattccttcctcctttctatc | RV primer for ptb21-mKate | Primer |
| HR544 | cggcaggtgtaagcggcgagccgcctctgtgccccagtttgctagggagg | RV primer for mKate2-wag31, mKate2 piece | Primer |
| HR545 | gcaaactggggcacagaggcggctcgccgcttacacctgccgacgtccac | FW primer for mKate2-wag31, wag31 piece | Primer |
| CMG154 | ccgcttacacctgccgacg | FW primer for wag31 | Primer |
| CMG61 | gtgctcaccaccggcctgacc | LamA knockout sequencing FW primer | Primer |
| CMG66 | caccaggccggccaggccgtag | LamA knockout sequencing RV primer | Primer |
| YL94 | ccatcgcgctctacctgctgctgac | Amplification of QcrB C-terminal upstream 500bp FW | Primer |
| YL95 | accttttccagaaccgtgatgaccgttggtctccccgttgccg | Amplification of QcrB C-terminal upstream 500bp RV | Primer |
| YL97 | accaacggtcatcacggttctggaaaaggtgaagaactg | Amplification of msfGFP FW | Primer |
| YL99 | ctccaattcgccctattatttgtagagttcatccatgccgt | Amplification of msfGFP RV | Primer |

| | | | |
|---|---|---|---|
| YL101 | gaactctacaaataatagggcgaattggagctccac | Amplification of HygR + loxP FW | Primer |
| YL103 | cgagtatctcagcgatacttctagactcgaggtaccggcg | Amplification of $Hyg^R$ + loxP RV | Primer |
| YL104 | tcgagtctagaagtatcgctgagatactcggatcgccgcaattcctc | Amplification of QcrB C-terminal downstream 500bp FW | Primer |
| YL105 | tgtcagctccgcgacacgccgct | Amplification of QcrB C-terminal downstream 500bp RV | Primer |
| KG7 | gcttaattaagaaggagatatacatatgaagacggtcaaggtcaacatcacg | FW primer for mycoQUEEN2m | Primer |
| KG8 | ctagggtccccaattaattagctaaagcttttacttcatctcggcgacc | RV primer for mycoQUEEN2m | Primer |
| HR242 | attaagaaggagatatacatatggcagagttgacaatctcggct | FW primer for AtpA-msfGFP, AtpA piece | Primer |
| HR243 | ctttactcgagccgccggccttcttgggtgccgg | RV primer for AtpA-msfGFP, AtpA piece | Primer |
| HR244 | aagaaggccggcggctcgagtaaaggtgaagaactgttc | FW primer for AtpA-msfGFP, msfGFP piece | Primer |
| HR245 | attaagaaggagatatacatatggcagccacactgcgcgaacta | FW primer for AtpG-msfGFP, AtpG piece | Primer |
| HR246 | ctttactcgagccgcctttcgagccggccagcgc | RV primer for AtpG-msfGFP, AtpG piece | Primer |
| HR247 | ggctcgaaaggcggctcgagtaaaggtgaagaactgttc | FW primer for AtpG-msfGFP, msfGFP piece | Primer |
| HRCT1 | gctagttaactacgtcgacatcga | Universal backbone RV primer | Primer |
| HRCT2 | ggctctgggagtacccgtg | Universal backbone FW primer | Primer |

| | | | |
|---|---|---|---|
| Y50A oligo | ccagcggtgtactgttcggcctccggcgcggaggccgcctgggagtagaactcggtctcaccggtctccg | Making $LamA_{Y50A}$ variant at the native locus | Oligo |
| $Hyg^{repair}$ | cggtccagcagccggggcgagaggtagccccacccgcggtggtcctcgacggtcgccgcg | Repair the mutated hygromycin cassette for SNP recombineering | Oligo |
| AtpA truncation ultramer | tgcggaccttgacggattccttctccaggtcctcgggatcgagggcctcggcgttctcggagacgaccacggtttgtaccgtacaccactgagaccgcggtggttgaccagacaaacccgagctgccgtcagaggcctggaagcccttcttgaattcgttgatgaccgagaccagcttctcctcggct | For ORBIT mediated truncation of AtpA C-terminal region | Ultramer |

### Citations

1. M. J. Smeulders, J. Keer, R. A. Speight, H. D. Williams, Adaptation of <i>Mycobacterium smegmatis</i> to Stationary Phase. *Journal of Bacteriology* **181**, 270-283 (1999).
2. J. J. Baker, R. B. Abramovitch, Genetic and metabolic regulation of Mycobacterium tuberculosis acid growth arrest. *Scientific Reports* **8**, (2018).
3. W. Lee, B. C. Vanderven, R. J. Fahey, D. G. Russell, Intracellular Mycobacterium tuberculosis Exploits Host-derived Fatty Acids to Limit Metabolic Stress. *Journal of Biological Chemistry* **288**, 6788-6800 (2013).
4. H. Yaginuma *et al.*, Diversity in ATP concentrations in a single bacterial cell population revealed by quantitative single-cell imaging. *Scientific Reports* **4**, 6522 (2014).
5. M. C. Martini, Y. Zhou, H. Sun, S. S. Shell, Defining the Transcriptional and Post-transcriptional Landscapes of Mycobacterium smegmatis in Aerobic Growth and Hypoxia. *Frontiers in Microbiology* **10**, (2019).
6. K. C. Murphy, K. Papavinasasundaram, C. M. Sassetti. (Springer New York, 2015), pp. 177-199.
7. K. C. Murphy *et al.*, ORBIT: a New Paradigm for Genetic Engineering of Mycobacterial Chromosomes. *mBio* **9**, (2018).
8. K. C. Murphy, in *Mycobacteria Protocols,* T. Parish, A. Kumar, Eds. (Springer US, New York, NY, 2021), pp. 301-321.
9. S. L. Tran, G. M. Cook, The F1 Fo-ATP Synthase of Mycobacterium smegmatis Is Essential for Growth. *Journal of Bacteriology* **187**, 5023-5028 (2005).
10. J. Schindelin *et al.*, Fiji: an open-source platform for biological-image analysis. *Nature Methods* **9**, 676-682 (2012).
11. S. Berg *et al.*, ilastik: interactive machine learning for (bio)image analysis. *Nature Methods* **16**, 1226-1232 (2019).
12. T. Falk *et al.*, U-Net: deep learning for cell counting, detection, and morphometry. *Nature Methods* **16**, 67-70 (2019).
13. L. Shao, P. Kner, E. H. Rego, M. G. L. Gustafsson, Super-resolution 3D microscopy of live whole cells using structured illumination. *Nature Methods* **8**, 1044-1046 (2011).
14. R. Fiolka, L. Shao, E. H. Rego, M. W. Davidson, M. G. L. Gustafsson, Time-lapse two-color 3D imaging of live cells with doubled resolution using structured illumination. *Proceedings of the National Academy of Sciences* **109**, 5311-5315 (2012).
15. R. Förster *et al.*, Simple structured illumination microscope setup with high acquisition speed by using a spatial light modulator. *Opt. Express* **22**, 20663-20677 (2014).
16. J. M. Belardinelli *et al.*, The MmpL3 interactome reveals a complex crosstalk between cell envelope biosynthesis and cell elongation and division in mycobacteria. *Scientific Reports* **9**, (2019).
